# Blocking the VLA4/VCAM1 axis prevents infarct-induced neurodegeneration and promotes vascular integrity

**DOI:** 10.1101/2025.06.25.661593

**Authors:** Kristy A. Zera, Karen Bradshaw, Li Zhu, Oliver Hahn, Aulden Foltz, Todd Peterson, Hanadie Yousef, Davis Lee, Elizabeth Mayne, Nima Aghaeepour, Tony Wyss-Coray, Marion S. Buckwalter

**Affiliations:** Department of Neurology and Neurological Sciences, Stanford University School of Medicine, Stanford, CA, USA; VA Palo Alto Health Care System, Palo Alto, CA, USA; Department of Anesthesiology, Perioperative & Pain Medicine, Stanford University School of Medicine, Stanford, CA, USA; Department of Biomedical Data Sciences, Stanford University School of Medicine, Stanford, CA, USA; Department of Pediatrics, Stanford University School of Medicine, Stanford, CA, USA; Department of Neurosurgery, Stanford University School of Medicine, Stanford University, Stanford, CA, USA

**Keywords:** Stroke, cognitive decline, VCAM1, pericyte, blood-brain barrier, vascular dysfunction

## Abstract

Infarct-induced neurodegeneration occurs chronically after stroke, doubling the risk of dementia. Endothelial vascular cell adhesion molecule 1 (VCAM1) facilitates blood-brain barrier opening and immune cell diapedesis by binding very late antigen 4 (VLA4) on immune cells. We hypothesized that vascular dysfunction persists after stroke and contributes to chronic neuroinflammation and cognitive decline via signaling through the VLA4/VCAM1 axis.

We used adult (3-5 month old) and middle-aged (10 month old) C57BL/6J male & female mice and a permanent middle cerebral artery occlusion stroke model. Sham surgery consisted of an identical procedure without occlusion of the artery. We quantified vascular integrity using blood vessel length, pericyte coverage of vasculature, tight junctions, and extravascular fibrinogen leakage by immunostaining. Cognitive testing was performed using both Barnes maze and novel object recognition prior to stroke, and 1 and 6 weeks after stroke, and replicated in both male and female mice. We utilized anti-VCAM1, anti-VLA4 or isotype control antibodies to block VCAM1 or VLA4 function, and then to confirm mechanisms we utilized single cell RNA sequencing on immune and endothelial cells, aptamer-based plasma proteomics, and additional immunostaining for vascular integrity.

Mouse brains exhibited signs of persistent vascular dysfunction and loss of blood-brain barrier integrity at 8 weeks after stroke, compared to sham animals. We observed reduced ZO-1 tight junction and pericyte coverage of vasculature, and increased extravascular fibrinogen. Mice with stroke also developed a cognitive deficit in both Barnes maze and novel object by 6 weeks. Treatment with anti-VCAM1 or anti-VLA4 resulted in mice with stroke performing as well as sham mice treated with isotype control antibody on both the Barnes maze and novel object recognition tasks. Anti-VCAM1 and anti-VLA4 both increased expression of blood-brain barrier maintenance genes in brain endothelial cells, while only minimally altering immune cell gene expression. Immune cell infiltration was reduced by anti-VCAM1 but not anti-VLA4 in tissue sections. In contrast to this, both antibodies increased blood vessel length and pericyte vascular coverage. Finally, extravascular fibrinogen was reduced by both antibody treatments in multiple brain regions.

Together, our findings establish the VLA4/VCAM1 axis as a promising target to preserve vascular integrity and prevent cognitive decline late after stroke. Our data is consistent with a model where blocking either VCAM1 or VLA4 chronically after stroke promotes new blood vessel growth and maturation and restores the blood-brain barrier to prevent infarct-induced neurodegeneration.

## Introduction

Cognitive dysfunction is a common and debilitating complication of stroke.^1^ Stroke occurs in over 12.2 million people worldwide each year, and independently doubles the risk of subsequent cognitive impairment for at least a decade.^1–4^ This is irrespective of established vascular dementia risk factors and is not prevented by reducing stroke incidence. Indeed, cognitive decline affects up to one-third of stroke survivors within five years, and represents a significant impediment to their quality of life.^1,5,6^ A survey of over 1200 stroke survivors found that almost 60% identify problems with concentration and memory as their primary unmet therapeutic need.^7^ Healthcare providers and caregivers of stroke survivors have also identified an understanding of cognitive changes after stroke as the foremost research priority in the field.^8^

One key mechanistic insight into cognitive decline after stroke is that it is biphasic. Acutely, the initial cognitive impairment is directly related to the location and severity of the initial infarct.^9,10^ After this initial phase, people with stroke remain at risk of delayed cognitive decline. We have coined the term “infarct-induced neurodegeneration” to describe this delayed risk. Infarct-induced neurodegeneration comprises approximately two thirds of the ten-year dementia risk, it is not mitigated by preventing additional strokes, and it is not associated with stroke location or severity.^1,11,12^ Importantly, because infarct-induced neurodegeneration is delayed, we are focused on understanding its mechanism in order to develop treatments to mitigate post-stroke dementia.

We first developed a mouse model to understand how infarct-induced neurodegeneration occurs, using a focal cortical stroke in wildtype mice.^13^ Mice exhibit no acute cognitive effect from the stroke, which is critical to allow us to separate lesion effects from the effects of the delayed neurodegenerative process. Then, they develop a cognitive deficit within 7 weeks, progressive loss of long-term potentiation over 14 weeks, and chronic immune infiltrates in the stroke scar. Notably, in this model the cognitive decline requires B lymphocytes, and we have also found B lymphocytes in the brains of humans who die with stroke and dementia.^13^ Together, these findings highlight an important role for chronic neuroinflammation in the development of infarct-induced neurodegeneration. However, humans do not have consistently high numbers of B lymphocytes at death, so we speculate that other mechanisms may be important for perpetuation of infarct-induced neurodegeneration.

Here, we investigated blood-brain barrier opening as a potential mechanism that might persist chronically and contribute to infarct-induced neurodegeneration. Immune cell trafficking across the blood-brain barrier is a tightly regulated process, which requires both bi-directional signaling and physical interaction between immune and endothelial cells. Immediately after a stroke, innate immune cells traffic to the brain through a leaky and disrupted blood-brain barrier.^14,15^ A leaky blood-brain barrier could itself contribute to dementia risk directly through admission of neurotoxic plasma proteins to the brain, as well as by promoting neuroinflammation.^16–18^ Therefore, we hypothesized that cerebrovasculature remains leaky chronically after stroke, and that this dysfunctional vasculature contributes to delayed dementia risk and infarct-induced neurodegeneration.

We first investigated vascular integrity and blood-brain barrier composition in the chronic phase after stroke, when mice have developed infarct-induced neurodegeneration and subsequent cognitive deficits in this model.^13^ We looked at key structural and functional aspects of the blood-brain barrier including pericyte coverage of vasculature, tight junction changes and fibrinogen leakage.

We next tested whether blocking the very late antigen-4/vascular cell adhesion molecule-1 (VLA4/VCAM1) axis would preserve vascular integrity and prevent infarct-induced neurodegeneration in mice. We chose to target this axis for a number of reasons. The VLA4 integrin receptor on immune cells binds to endothelial VCAM1 to initiate immune cell trafficking across the blood-brain barrier.^19,20^ It is a key signaling pathway by which immune cells, including B lymphocytes, enter tissues during inflammation. Additionally, VCAM1 expression increases in people with both vascular dementia and stroke, suggesting a potential causative link to dementia after stroke.^21,22^ Expression of VCAM1 is not only increased on endothelial cells after stroke, it is also increased in aging, where blocking antibodies improve cognition in aging mice, likely through a vasculoprotective mechanism.^23^ Thus, we hypothesized that blocking the VLA4/VCAM1 axis might improve cognition via both immune-dependent and vascular-dependent mechanisms in our mouse model of infarct-induced neurodegeneration. To test this, we treated middle-aged male and female mice with either anti-VCAM1 or anti-VLA4 antibodies, and used Barnes maze and novel object recognition to test cognitive outcomes. In addition, we investigated underlying mechanisms using immunostaining, plasma proteomics and single-cell RNA sequencing. Our findings support a critical role for the VLA4/VCAM1 axis in the development of infarct-induced neurodegeneration, and identify this axis as a potential therapeutic target to prevent cognitive decline after stroke.

## Methods

### Mice

Adult (3-5 month old) male and middle-aged (10 month old) male and female C57BL/6J mice were purchased from Jackson Laboratory (#000664). Animals were group housed 3-5 mice per cage, and allowed to acclimate for at least 1 week prior to experimentation. All experiments were performed in accordance with protocols approved by the Stanford University Institutional Animal Care and Use Committees (IACUC).

### Stroke surgeries

Stroke surgeries were performed as previously described.^24^ Briefly, mice were anesthetized with ∼2% isoflurane in 100% oxygen and maintained at 37°C with a temperature-controlled heating pad for the duration of the surgery. The incision site was cleaned with chlorhexidine (Henry Schein #5680808), and an incision was made along the skull. Next, the temporalis muscle was transected so that the middle cerebral artery (MCA) was visible through the skull. A craniotomy was drilled directly above the MCA, the meninges were carefully removed, and the MCA was permanently cauterized. The muscle and skin were replaced, and the incision was closed with surgi-lock glue (Meridian #78657941). Sham mice underwent the same procedure, including removal of meninges, but did not undergo MCA cauterization. Mice subsequently received a single subcutaneous dose of cefazolin (25 mg/kg; VWR #89149-888) and buprenorphine SR (1 mg/kg; Henry Schein #6030894) as antibiotics and analgesia, respectively. Middle-aged (10 month old) mice were then kept in a warmed cage until they were awake and mobile, approximately 30 minutes following the surgery. Adult (3 month old) sham and stroke mice were allowed to recover for 5 mins before spending 60 min in a temperature controlled 37°C chamber in 8% oxygen/92% nitrogen (dMCAO + hypoxia stroke). The purpose of the hypoxia was to increase infarct size and reduce variability between mice, although this step significantly increases mortality in older mice and was therefore not utilized.^24^

### Antibody Treatment

Adult (3 month old) and middle-aged (10 month old) C57BL/6J male and female mice received injections of anti-VCAM1 monoclonal antibody (α-VCAM1, BE0027, BioXCell), anti-VLA4 monoclonal antibody (α-VLA4, BE0071, BioXCell) or the appropriate isotype control antibody (α-IC, VCAM1: BE0088; VLA4: BE0090; BioXCell) at a dose of 9 mg per kg. For all treatment groups, the first dose at 4 hours was retro-orbital, and subsequent doses were intraperitoneal. Sham-α-IC and Stroke-α-IC mice were dosed with the appropriate isotype control antibody beginning 4 hours after stroke, and then every 3 days for the duration of the experiment, using both isotype control antibodies for experiments where treatment groups included both anti-VCAM1 and anti-VLA4 antibodies. Stroke-α-VCAM1 (Acute) and Stroke α-VLA4 (Acute) mice received a single dose of their respective antibody solution at 4 hours after surgery. They then received doses of isotype control antibody every 3 days for the duration of the experiment. Stroke-α-VCAM1 (Chronic) and Stroke-α-VLA-4 (Chronic) mice received one dose of isotype control antibody at 4 hours after stroke, and the first dose of anti-VCAM1 or anti-VLA4 solution 4 days after stroke. They received subsequent doses of anti-VCAM1 or anti-VLA4 every 3 days for the duration of the experiment. Thus, all mice received a total of 16 injections over the course of the experimental period.

### Cognitive Behavioral Testing

Mice were group housed and each cage contained a random mixture of mice from each treatment group. Male and female mice were tested in separate cohorts to avoid confounding scents. The order of testing, groups and start time of testing were maintained on each day to reduce circadian variability. Prior to the initiation of the experiment, mice were subjected to three days of handling. Prior to testing or handling on any given day, mice were acclimated to the testing room for at least 60 minutes.

#### Barnes Maze

A modified Barnes maze protocol was utilized as previously described.^23,25^ Briefly, a circular maze containing 16 holes around the outer edge was centered on a pedestal and elevated approximately 3 feet above the floor. All holes were open to the floor except for the escape hole, which consisted of a PVC elbow joint connector. Distinct visual cues were placed at four equally spaced points around the maze. An overhead light, two standing lights and a fan blowing on the maze provided motivation to learn the escape hole location, while also serving as supplemental visual cues. The escape hole position was aligned with one of the visual cues, and remained fixed for all trials. Mice performed four trials per day for four consecutive days. The starting location of the mouse was moved relative to the escape hole position at the start of every trial. Escape latency was defined as the time for a mouse to enter the escape hole. When a mouse failed to locate the escape hole within 90 seconds, it was gently guided towards the escape hole and assigned an escape latency of 90 seconds. The maze and escape holes were thoroughly cleaned with 10% EtOH after every trial and with 70% EtOH at the end of every testing day to eliminate olfactory cues. Testing was performed by an investigator blinded to experimental conditions.

#### Novel Object Recognition (NOR)

NOR was performed as previously described.^23,25^ Briefly, each cage of mice was habituated to an empty arena (40 × 40 × 35 cm^3^) containing wall-mounted visual cues for 5 min. Mice were then individually given one 5 min trial during which they explored two identical objects in fixed positions in the middle of the arena. Animals were randomly assigned to identical starting objects and new object pairs were utilized for each testing timepoint. The testing phase began 2 hours after completion of the first exploration phase. In the testing phase, mice explored the same arena for 5 min but with one object replaced by a novel, distinct object. The arena and objects were thoroughly cleaned with 10% EtOH to eliminate residual odors between mice and between testing sessions. Interactions with objects (sniffing or exploring within 2 cm of object or climbing on object; excluding time spent sitting on top of object) were manually timed in a blinded fashion. To assess performance on the task, a discrimination index (DI) score was calculated using the formula [(t_novel_-t_familiar_)/(t_novel_+t_familiar_)] to represent the proportion of time spent on the novel object. A score between 0 and -1 indicates a preference for the familiar object, a score between 0 and +1 indicates a preference for the novel object, and a score of 0 demonstrates no object preference.

### Mouse Brain Immunohistochemistry and Immunofluorescence

Immunohistochemistry staining was performed as previously described on PFA-fixed 40 μm coronal brain sections using 3,3’-diaminobenzidine visualization.^13,26^ Primary antibodies were against B220 (biotinylated, 1:500; BD Biosciences, 553085), CD3ε (1:500; BD Biosciences, 550277), and NeuN (1:500; Millipore, MAB377B). The specificity of primary antibody staining was confirmed using control sections treated with secondary antibody alone, and by visual inspection of expected cellular morphology. Images were taken using a Keyence BZ-X710 microscope. Quantification of immunostaining was performed on two consecutive sections per mouse, by a researcher blinded to the treatment group. Images were taken at 20x magnification, and quantification was performed on the full field of view. B220 immunostains were quantified using thresholding in ImageJ and reported as percent of the area covered by antibody staining. The total number of CD3+ T cells per field of view was also counted using ImageJ. For atrophy quantification, every 6th section was stained using NeuN with a cresyl violet counterstain and captured with a slide scanner (PrimeHisto XE Histology # 489780). ImageJ was used to manually trace the infarct core, ipsilateral hemisphere, and contralateral hemisphere. Total brain atrophy was calculated and reported as percent loss of volume compared to the contralateral hemisphere.

Immunofluorescent staining was also performed as previously described following standard staining techniques.^26^ Primary antibodies were used against CD13 (1:500; Bio-Rad, MCA2183GA), CD31 (1:200; Novus Biologicals, AF3628 or 1:300, BD Biosciences, 550274), fibrinogen (1:1000; DAKO, A0080), ZO-1 (1:100; ThermoFisher, 33-9100) and GFAP (1:1000; Abcam, Ab4674). To image, Z-stacks of 5 images each were taken at 40x magnification using a Leica TCS-SPE confocal microscope in the medial peri-infarct region of the cortex, defined as one field of view away from the stroke border and at least one view field above the corpus callosum, or in the stroke core. A GFAP co-stain was used to define the stroke border and imaging location. Two brain sections per mouse were used for analysis. Sections were excluded from the analysis if they showed irregular cellular morphology. Pericyte vascular coverage was quantified in the peri-infarct cortex using ImageJ by first defining vasculature by tracing the CD31 stain, and then estimating the percent of total CD31+ vessel surrounded or covered by CD13+ staining. If only one side of the vessel was closely associated with an adherent pericyte, this was counted as 50% vessel coverage. The average CD13+ vessel coverage was then calculated for each animal. Vessel number and length were then calculated from the trace of the CD31 stain. Extravascular fibrinogen was quantified with a custom written macro that created a mask of the CD31 stain and then quantified the fibrinogen stain outside of the vasculature using thresholding in images from the stroke core and peri-infarct cortex.

### Single Cell RNA Sequencing

#### Brain dissociation

Brain tissue was dissociated with the Miltenyi neural dissociation kit following the manufacturer’s instructions, with minor adjustments to optimize the isolation of endothelial and immune cells. In brief, 4–5 mice per group were anesthetized with ketamine/xylazine solution and perfused with 15mL ice cold PBS following cardiac blood collection. The stroke core and peri-infarct cortex, defined as ∼2mm surrounding the visible lesion, were dissected and pooled by condition, minced, and digested using the neural dissociation kit (Miltenyi, 130–092-628). Brain homogenates were further diluted in Dulbecco’s PBS (dPBS) and filtered through a 100 µm strainer. The filtered homogenate was then centrifuged (300 *g* for 10 minutes), and the pellets were resuspended in 0.9 M sucrose in dPBS followed by centrifugation for 20 minutes at 850 *g* at 4 °C to separate the myelin. Cell pellets were resuspended in FACS buffer (1% BSA in PBS with 2 mM EDTA) and blocked for 5 min on ice with Fc block (CD16/CD32, BD 553141), followed by staining for 10 minutes with anti-CD31-PE/TR594 (1:200, BD 563616) and anti-CD45-PE/Cy7 (1:200, BD 103114) antibodies. Dead cells were excluded by staining with SYTOX Blue Dead Cell Stain (1:5,000, ThermoFisher Scientific S34857).

#### Fluorescence activated cell sorting (FACS)

Brain endothelial and immune cells were sorted on the FACSAria 3.3 (BD Biosciences) system using the FACSDiva software (BD Biosciences). Cells were gated on forward scatter (FSC, size) and side scatter (SSC, internal structure). FSC-A and FSC-W plotting was used to discriminate single cells from cell doublets or aggregates. SYTOX Blue+ dead cells were also excluded via gating. CD31+CD45− cells were defined as the endothelial cell population, and CD31lowCD45+ cells were defined as the immune cell population. Cells were sorted into 0.1% BSA solution with 0.2U/µL RNAse inhibitor in PBS and then pelleted at 400 *g* for 5 minutes at 4 °C. They were then resuspended in 50 µL 0.1% BSA with 0.2U/µL RNAse inhibitor before processing for single-cell RNA-sequencing.

#### RNA Extraction, Library Preparation and RNA Sequencing

Reagents of the Chromium Single Cell 3’ GEM & Gel Bead Kit v3.1 (10X Genomics, 1000121) were thawed and prepared according to the manufacturer’s protocol. Cells and master mix solution were adjusted to target 10,000 cells per sample and loaded on a standard Chromium Controller (10X Genomics, 1000204) according to manufacturer’s protocols. We applied 11 PCR cycles to generate cDNA. Library construction was conducted using the Chromium Single Cell 3’ Library Construction Kit v3 (10X Genomics, 1000121). All reaction and quality control steps were carried out according to the manufacturer’s protocol and with recommended reagents, consumables, and instruments. We used 11 PCR cycles for library generation. Quality control of cDNA and libraries was conducted using a Bioanalyzer (Agilent) at the Stanford Protein and Nucleic Acid Facility. Illumina sequencing of the resulting libraries was performed by Novogene (https://en.novogene.com/) on an Illumina NovaSeq S4 (Illumina). Base calling, demultiplexing, and generation of FastQ files were conducted by Novogene.

#### Data processing & Analysis

Cell Ranger (v.6.0.0) analysis pipelines were utilized to align reads to mm10-GRCh38 hybrid reference genome and count barcodes/UMIs. Reads mapping to introns were not counted. Outliers with a high ratio of mitochondrial (more than 10%, fewer than 400 features) relative to endogenous RNAs and homotypic doublets (more than 6,000 features) were removed in Seurat. Cells were also removed if they expressed fewer than 100 unique genes. Genes not detected in any cell were removed from subsequent analysis. Log-normalized (scale factor 10,000 and pseudocount of 1) expression values of genes generated by the Seurat NormalizeData command were used for clustering and differential gene expression analysis.

Mitochondrial and cell cycle gene regression was performed to ensure that clustering reflects true cell type identities. Principal component analysis was performed on the 2000 most variable genes of the dataset followed by UMAP clustering using the Seurat R Package (v. 4.3.0.1) on the first 20 principal components.

Identification of significant clusters was performed via the standard FindClusters command in Seurat with a 1.0 resolution. CD31+ and CD45+ cells were sub-clustered based on *Pecam1* and *Ptprc* expression, respectively. CD31+ cells and CD45+ cells underwent principal component analysis and were re-clustered on a resolution parameter of 0.2 and 0.4, respectively. Cell counts were found once the clusters were subset by condition. To pass QC, cells had a minimum of 300 reads, and on average had 5594 reads and 2208 expressed genes per cell. The annotation of each cluster was manually determined using published marker genes for each cell subtype (Supplementary Table 1). Next, MAST differential gene expression analysis between conditions was performed using Mast (v. 1.24.1), and Gene Ontology pathway analysis was performed with Enrichr.^27–29^ Finally, the normalized average expression of each gene within each cluster was generated using the Dotplot function of Seurat and plotted with GraphPad Prism 9.

### Plasma Proteomics

#### Blood Collection and Processing

Blood was collected from the left ventricle using an EDTA-impregnated syringe, prior to perfusion for brain collection. Whole blood was deposited into Microvette 500 K3EDTA tubes (Sarstedt #201341), and tubes were centrifuged at 4°C for 15 minutes at 2,000*g*. The resulting supernatant was collected as plasma and stored at -80°C prior to use for proteomic analysis.

#### SomaLogic Proteomics Analysis

Plasma protein profiling was performed using the SomaLogic SOMAScan platform (SomaLogic Inc.). This technology employs slow off-rate modified DNA aptamers (SOMAmers) to quantify the relative abundance of 5,284 mouse proteins within plasma with high specificity. The platform’s performance characteristics have been previously described.^30,31^ Individual protein differences between Stroke-α-IC and Stroke-α-VCAM1 (Chronic) treatment groups were analyzed with a Mann Whitney test with *p* < 0.05 considered significant. Data were processed using GraphPad Prism 10 software. Statistical outliers were removed utilizing the ROUT method (*Q* = 0.1%). A univariate Spearman correlation was performed to analyze the relationship between protein concentration, regardless of treatment group, and cognitive function. To assess cognitive function, a cognitive score was calculated by averaging the Barnes maze performance from trials 3 & 4 of the final day of testing for each mouse in Figure 2B. Therefore, a lower cognitive score represents better cognitive function.

#### Statistics

Experimenters were blind to treatment status throughout data collection and analysis. Statistical testing was performed in GraphPad Prism 10 or R version 4.2.2. Behavioral data were analyzed with a two-way repeated measures ANOVA with Tukey’s post-hoc test. Immunostaining data were analyzed with a Student’s t test (2 groups) or one-way ANOVA with Tukey’s post-hoc test (3 or more groups). Plasma proteomic data were analyzed with a Mann Whitney (mouse) or Kolmogorov-Smirnov (human) test. For all statistical testing, *p* < 0.05 was considered statistically significant.

## Results

### Vascular dysfunction persists in chronic stroke

To investigate whether there is a chronically dysfunctional blood-brain barrier around the stroke lesion, we analyzed structural and functional measures of blood-brain barrier integrity at 8 weeks after stroke, which is after cognitive impairment develops. Pericytes are required to induce tight junctions between endothelial cells and to recruit astrocytic endfeet to reduce barrier permeability.^32–34^ Thus, we analyzed vascular pericyte coverage as a measure of blood vessel maturation and blood-brain barrier integrity. In the peri-infarct cortex, stroked mice exhibited 36% less CD13+ pericyte coverage of CD31+ blood vessels than animals who underwent sham surgery (Figure 1A). We next analyzed ZO-1 immunostaining to assess tight junctions themselves. The relative coverage of ZO-1 immunostaining along vasculature in mice with stroke was reduced by 50% compared to their respective sham controls (Figure 1B). Finally, we formally quantified blood-brain barrier integrity via extravascular fibrinogen leakage into the brain parenchyma. In mice with stroke, extravascular fibrinogen covered 9.2% of the infarct core and 0.3% in the peri-infarct cortex, with no detectable leakage in sham animals (Figure 1C). Together, these data suggest that chronic lymphocyte trafficking into the brain after stroke is associated with persistent vascular dysfunction that does not resolve in the weeks after stroke.

**Figure 1.**
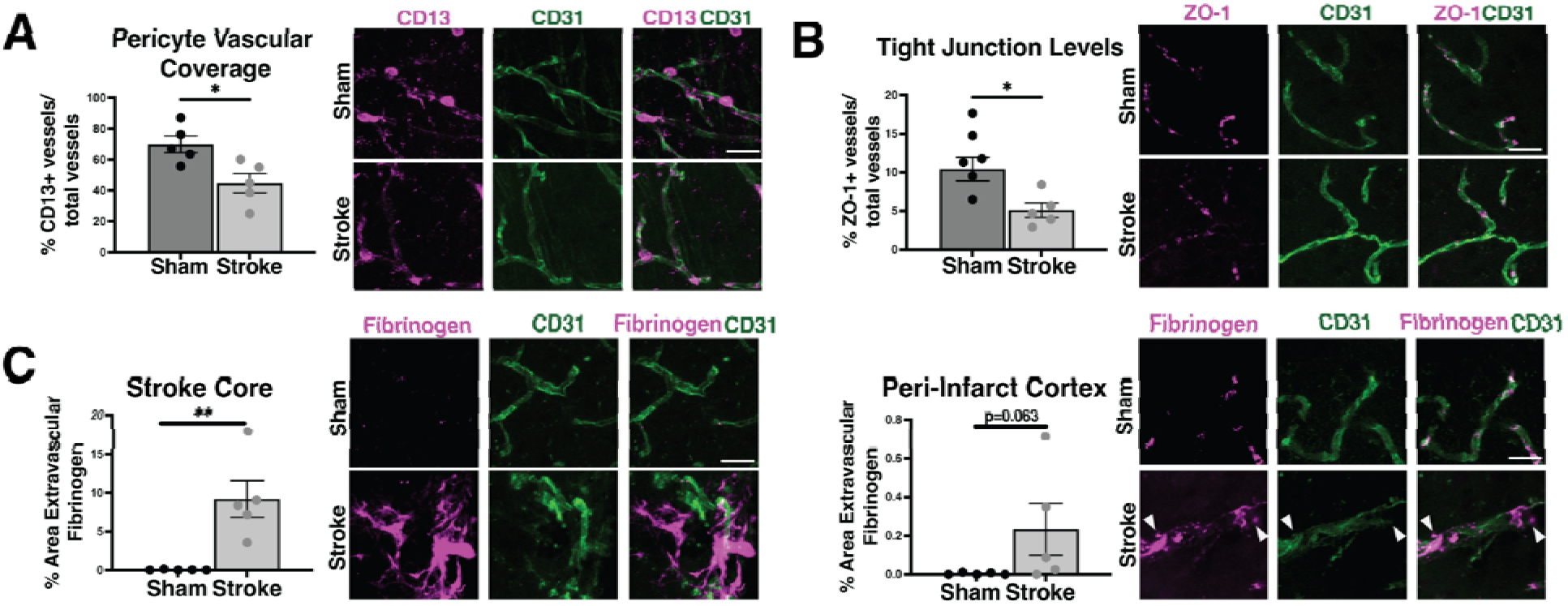
Vascular dysfunction persists chronically after stroke. A) Representative confocal images and quantification of pericyte (CD13+) coverage of CD31+ vasculature in the peri-infarct cortex of sham or stroked mice, 8 weeks after surgery. B) Representative confocal images and quantification of vascular (CD31+) tight junctions (ZO-1) in the peri-infarct cortex. C) Representative confocal images and quantification of the percent area covered by extravascular fibrinogen in the stroke core (left) and peri-infarct cortex (right) of the same mice. Scale bar, 20uM; Statistics, Student’s t-test Error bars, mean ± SEM; *p<0.05; **p<0.01.

### Chronic anti-VCAM1 prevents cognitive decline

Since VCAM1 is elevated after stroke and persists in dementia,^21,35–38^ and also promotes vascular leakiness and immune cell trafficking into the brain, we next tested its role in infarct-induced neurodegeneration. We first assessed the impact of acute versus chronic blockade with a VCAM1 blocking antibody (α-VCAM1) in 10-month-old C57BL/6J mice (Figure 2) to test whether blocking initial immune cell influx would ameliorate infarct-induced neurodegeneration or whether prolonged chronic blockade was needed. In these two paradigms, the acute treatment at 4 hours after stroke should largely affect innate immune cell trafficking while the chronic treatment starting 4 days after stroke should largely target adaptive immune cells. Both should affect vascular permeability. As expected for this mouse model, mice that received isotype control antibody (α-IC) perform normally on the Barnes maze and novel object recognition tasks before and 1 week after stroke, then exhibit delayed cognitive decline 6 weeks after stroke (Figure 2B-E). In the acute paradigm, Stroke-α-VCAM1 Acute, there was no effect on late cognitive decline after stroke in male mice (Figure 2B, D). In contrast, male mice treated chronically with α-VCAM1 (first at 4 days, then every 3 days; Stroke-α-VCAM1 Chronic) did not develop a cognitive deficit after stroke in either task (Figure 2B, D). This was also true when the experiment was repeated in female mice (Figure 2C, E). Thus, chronic, but not acute, treatment with α-VCAM1 prevented delayed cognitive decline after stroke in both sexes.

**Figure 2.**
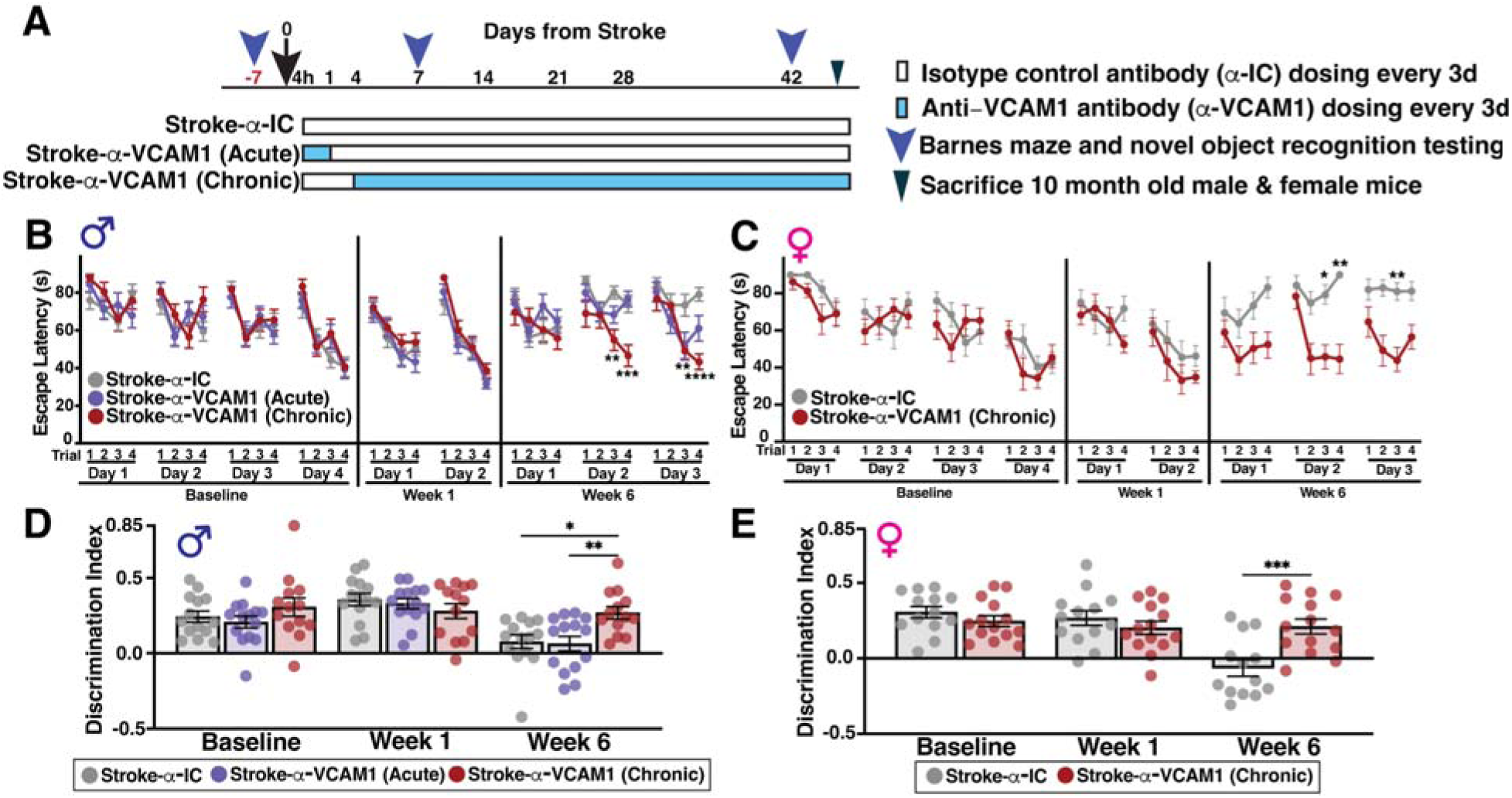
Chronic anti-VCAM1 antibody treatment prevents cognitive decline and reduces lymphocyte trafficking in male and female middle-aged mice. A) Diagram depicting the study design. All animals received doses at 4 hours and then every 3 days with isotype control antibody (α-IC) or VCAM1-blocking antibody (α-VCAM1) as depicted. B) Escape latency on the Barnes maze in male or (C) female mice (10 month old) treated with the appropriate isotype control antibody throughout (α-IC, n=14); a single anti-VCAM1 dose at 4 hours (Stroke-α-VCAM1 Acute; n=14) or anti-VCAM1 throughout (Stroke-α-VCAM1 Chronic, n=13). D) Discrimination index on the novel object recognition task in male or (E) female mice (10 month old) treated with the appropriate isotype control antibody throughout (α-IC, n=14); a single anti-VCAM1 dose at 4 hours (Stroke-α-VCAM1 Acute; n=14) or anti-VCAM1 throughout (Stroke-α-VCAM1 Chronic, n=13). Statistics, 2-way repeated measures ANOVA with Bonferroni’s post-hoc test (B-C) or one-way ANOVA with Tukey’s post hoc test (D-E). Error bars, mean ± SEM; *p<0.05; **p<0.01; ***p<0.001; ****p<0.0001.

There was no difference in brain atrophy chronically after stroke in both male and female mice after either acute or chronic α-VCAM1 administration (Supplementary Figure 1). Thus, the protective effects of α-VCAM1 treatment are not mediated by reducing stroke size or subsequent atrophy.

### Chronic anti-VLA4 treatment prevents cognitive decline after stroke

We next asked whether blocking VLA4, the binding partner of VCAM1 on immune cells, would be beneficial and whether it would be more or less effective at treating infarct-induced neurodegeneration than targeting VCAM1. As expected, middle-aged, 10 month old, male mice treated with an isotype control antibody (Stroke-α-IC) developed a robust cognitive deficit in both the Barnes maze and novel object recognition tasks at 6 weeks after stroke, compared to Sham-α-IC mice (Figure 3). This was also replicated with an independent cohort of female mice (Supplementary Figure 2). Cognitive decline was prevented equally in α-VCAM1 Chronic and α-VLA4 Chronic treatment groups on both the Barnes maze and novel object recognition tasks in both male and female mice (Figure 3, Supplementary Figure 2). Moreover, both α-VCAM1 and α-VLA4 treatment maintained the cognitive performance of mice with chronic stroke equivalent to that of sham animals at all timepoints, and on both cognitive tasks, and in mice of both sexes.

**Figure 3.**
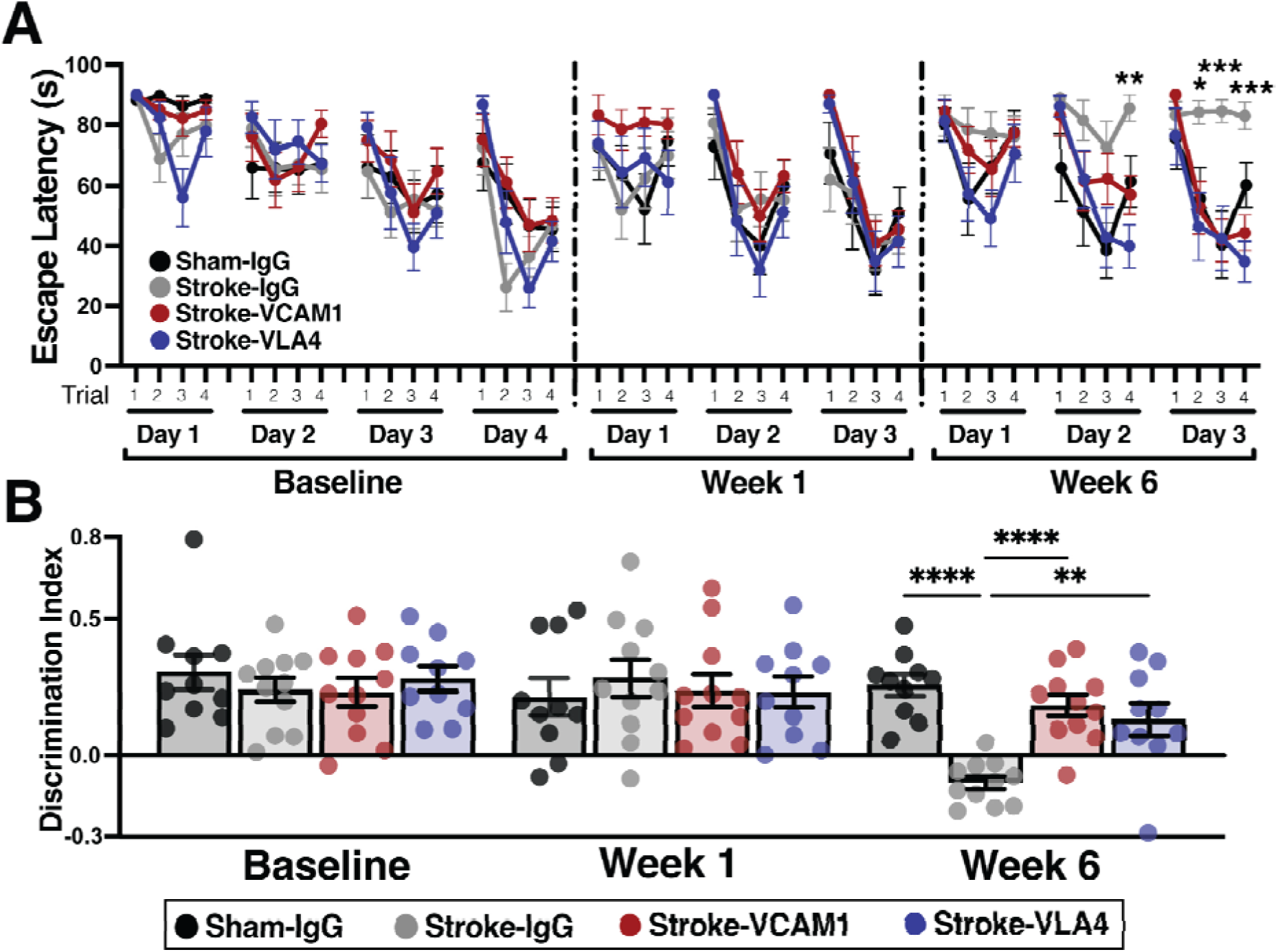
Both anti-VCAM1 and anti-VLA4 chronic antibody treatment prevent cognitive decline in middle-aged mice. A) Escape latency on the Barnes maze and B) discrimination index on the novel object recognition task for 10 month old male mice treated with isotype control antibody (α-IC; n=10 sham and 11 stroke), anti-VCAM1 (α-VCAM1; n=11) or anti-VLA4 (α-VLA4; n=10) chronically. Statistics, 2-way repeated measures ANOVA with Bonferroni’s post-hoc test; Error bars, mean ± SEM; *p<0.05; **p<0.01; ***p<0.001; ****p<0.0001.

As with α-VCAM1 treatment in middle-aged (10 month old) mice, there was no effect on brain atrophy in adult (3 mo) mice after α-VCAM1 or α-VLA4 chronic treatment compared to stroked mice treated with isotype control antibody (Stroke-α-IC; Supplementary Figure 3).

### Both anti-VCAM1 and anti-VLA4 had only modest effects on immune cells in chronic stroke

Since the VLA4/VCAM1 axis impacts both endothelial cell activation and immune cell trafficking, we next performed single cell RNA sequencing of both brain immune (CD45+) and endothelial (CD31+) cells. We isolated cells from the stroke scar and peri-infarct cortex of mice from all four treatment groups in Figure 3, ten weeks after surgery. For all groups, treatment commenced 4 days after surgery and continued every 3 days until the end of the experiments. This included mice after stroke or sham surgery treated with isotype control antibody, and mice with stroke treated chronically with either α-VCAM1 or α-VLA4. We analyzed 15,263 total cells (7,942 CD45+ and 7,321 CD31+). Cells were clustered with the UMAP algorithm to reduce dimensionality, and cell-specific markers were used to annotate 11 CD45+ clusters, comprising all major peripheral and brain-resident immune cell types (Figure 4A; Supplementary Figure 4B). Similarly, we identified 5 CD31+ cell clusters; arterial, 2 capillary and 2 venous endothelial cell populations (Figure 4B; Supplementary Figure 4A).

**Figure 4.**
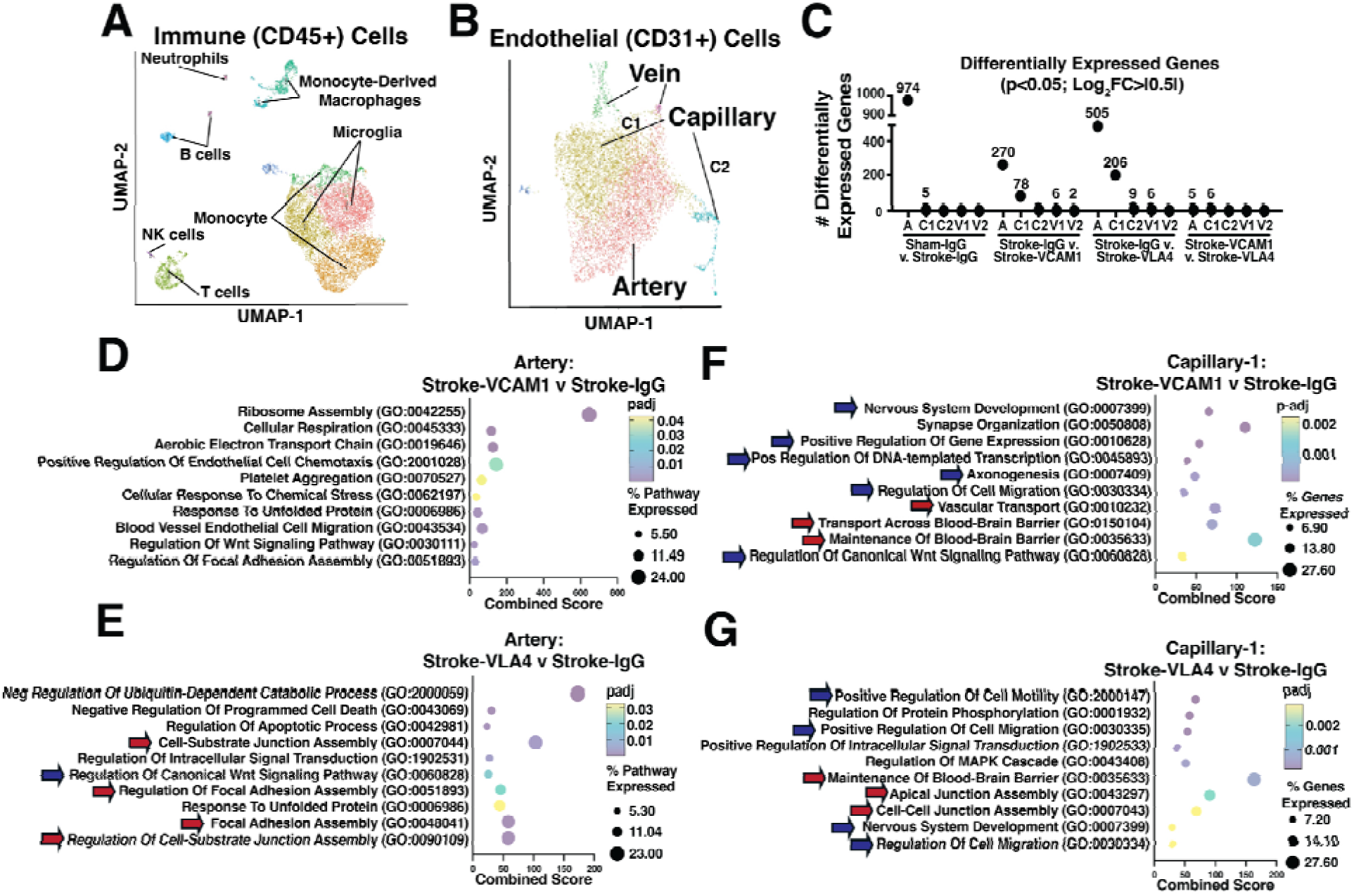
Anti-VCAM1 and anti-VLA4 treatment induces changes in endothelial cells which reflect generation of new and more mature blood vessels. Unbiased UMAP visualization of (A) immune and (B) endothelial cell populations isolated from the stroke scar of 10 month old male mice, 10 weeks after stroke, based on single cell RNA sequencing. Each sample represents a pool of 4-5 mice in 4 groups, sham and stroke mice treated with isotype control antibody (Sham-α-IC and Stroke-α-IC), and stroke mice treated chronically with anti-VCAM1 (Stroke-α-VCAM1) or anti-VLA4 (Strokeα-VLA4) antibody. For all groups, treatment was every 3 days, beginning 4 days after surgery for a total of 10 weeks. C) A graphical representation of the number of differentially expressed genes in each subpopulation of endothelial cells, separated by the specific comparisons of interest. D-G) Over-representation pathway analysis on the differentially expressed genes in the artery and capillary-1 clusters. Comparisons are between Stroke-α-IC and either Stroke-α-VCAM1 (D,F) or Stroke-α-VLA4 (E,G) treatment groups. The top pathways are included here, along with the combined score (x-axis), the percent of genes expressed within each pathway (dot size) and p value of pathway enrichment (dot color). Blue arrows identify pathways involved in vessel growth, while red arrows identify pathways involved in vessel maturation and blood-brain barrier maintenance.

As a fraction of total cells counted, the proportion of peripheral immune cells in the brain increased after stroke. The proportion of T cells increased from 6.2% in Sham-α-IC to 11.2% in Stroke-α-IC, and the proportion of B cells also increased from 1.9% to 5.2% in the same groups (Supplementary Figure 4C). Chronic α-VCAM1 or α-VLA4 treatment then reduced proportions of T and B lymphocytes down to that seen in the Sham-α-IC group. In chronic α-VCAM1 treated mice with stroke (Stroke-α-VCAM1 Chronic), the proportion of T cells was 5.1% and B cells 1.4% (Supplementary Figure 4C). In chronic α-VLA4 treated mice with stroke (Stroke-α-VLA4 Chronic), the proportions were 7.4% and 1.7%, respectively. For CD31-expressing vascular cells, after stroke there was a higher proportion of cells within the two capillary clusters, and fewer cells in the arterial cell cluster. Stroke increased the proportion of capillary endothelial cells from 32.5% in Sham-α-IC to 42.3% in Stroke-α-IC, and reduced the proportion of arterial endothelial cells from 62.2% in Sham-α-IC to 53.5% in the Stroke-α-IC sample. Both α-VCAM1 and α-VLA4 chronic treatment also slightly increased the proportion of capillary endothelial cells and reduced the proportion of arterial endothelial cells after stroke (Supplementary Figure 4D).

Despite the increase in proportion of peripheral immune cells in the brains of mice with chronic stroke, there were very few significant (*p*<0.05; Log_2_FoldChange>|0.5|) transcriptomic changes in immune cells, compared to cells from the Sham-α-IC sample (Supplementary Figure 5). A similar pattern was observed after α-VCAM1 or α-VLA4 treatment, compared to the Stroke-α-IC group. In fact, other than the Monocyte-1 cluster (Figure 4A, bottom cluster) which had 94 differentially expressed genes when comparing Stroke-α-VLA4 to Stroke-α-IC, no other bioinformatic comparison of peripheral immune cell types found more than 19 differentially expressed genes.

We next examined T and B lymphocyte accumulation in the brain by immunohistochemistry (Supplementary Figure 6). As with cognition, acute treatment with α-VCAM1 had no effect on either T or B lymphocyte accumulation in the brain, while chronic treatment did partially reduce lymphocytes. It reduced the number of CD3+ T cells in the infarct core by 44% in male and 41% in female mice (Supplementary Figure 6A, B). Similarly, chronic α-VCAM1 treatment reduced the percent area covered by B lymphocytes in the stroke core by 53% in male and 62% in female mice (Supplementary Figure 6C,D). Thus, a reduction in adaptive immune cells but not in brain atrophy is associated with the prevention of cognitive decline after stroke following chronic α-VCAM1 treatment.

Comparing the effects of chronic α-VCAM1 to chronic α-VLA4 revealed that α-VCAM1 again significantly reduced T lymphocyte accumulation in the stroke core by 41% (Supplementary Figure 7A) and B lymphocyte accumulation by 83% (Supplementary Figure 7C). In contrast, chronic α-VLA4 did not change T or B cell accumulation in the infarct core (Supplementary Figure 7B,D), despite the fact that it had equivalent effects on cognition as α-VCAM1 treatment. Similarly, and consistent with our findings with α-VCAM1, a single acute dose of either α-VCAM1 or α-VLA4 (Stroke-α-VCAM1 Acute or Stroke-α-VLA4 Acute, respectively) did not affect T or B lymphocyte accumulation in the infarct core (Supplementary Figure 7).

### Both anti-VCAM1 and anti-VLA4 promote vessel growth & maturation

In contrast to the less impressive results with immune cells, both blocking antibodies induced dramatic changes in gene expression in endothelial cells with stroke alone (Sham-α-IC vs. Stroke-α-IC), and also when α-VCAM1 or α-VLA4 treatment was compared to Stroke-α-IC. When using a differentially expressed gene (DEG) cutoff of *p*<0.05 and Log_2_FoldChange>|0.5| within each subpopulation of endothelial cells, arterial cells were the most changed by stroke, with increased expression of 501 genes and decreased expression of 473 genes (Figure 4C). Additionally, chronic treatment with α-VCAM1 or α-VLA4 primarily affected the arterial and capillary-1 (left cluster, Figure 4B) endothelial cell clusters, which were also the two clusters with the largest proportional population changes after stroke, or either antibody treatment. After α-VCAM1 treatment, arterial endothelial cells increased expression of 200 genes and decreased expression of 70 genes, while capillary-1 endothelial cells increased expression of 50 and decreased expression of 28 genes, compared to Stroke-α-IC cells. Similarly, after chronic α-VLA4 treatment, arterial cells increased expression of 268 genes and decreased expression of 237 genes, while capillary-1 cells increased expression of 120 and decreased expression of 86 genes. In contrast to the large number of gene expression changes when comparing antibody treatment to the Stroke-α-IC group, the differences between the α-VCAM1 and α-VLA4 treated groups were minimal. Specifically, there were <10 differentially expressed genes when directly comparing α-VCAM1 to α-VLA4 treated endothelial cells from any cluster (Figure 4C).

We next used over-representation analysis to understand the biological pathways underpinning the transcriptomic changes induced by stroke, and chronic α-VCAM1 or α-VLA4 antibody treatment after stroke. We focused on the arterial and capillary-1 clusters because they had the highest number of changed genes. In the arterial cell cluster, stroke reduced expression of genes in pathways involving cell proliferation and development, and new vessel growth. In contrast, both α-VCAM1 and α-VLA4 treatment enriched multiple pathways related to cell proliferation and migration, pointing to angiogenesis and new vessel growth after chronic antibody treatment (Figure 4D,E, blue arrows). Both antibody treatments also enriched multiple pathways involved in vessel maturation compared to the Stroke-α-IC group in the arterial cell cluster (Figure 4D,E, red arrows). Consistent with abnormalities in vascular growth and maturation in chronic stroke, many of the same genes involved in cellular proliferation pathways were also reduced in the capillary-1 cluster in the Stroke-α-IC group compared to Sham-α-IC. After both chronic α-VCAM1 and α-VLA4 treatment, many of the same pathways were enriched in the capillary-1 cluster, compared to the Stroke-α-IC condition (Figure 4F, G).

Indeed, key genes involved in cell-cell junction formation, pericyte recruitment and cytoskeleton assembly, all of which are critical for vessel growth and maturation, all followed a consistent pattern. Overall, stroke reduced expression of genes related to vascular growth and maturation, and also reduced the proportion of endothelial cells expressing those genes (left panel, Figure 5A). In contrast, chronic treatment of mice after stroke with either VCAM1 or VLA4 blocking antibodies reversed this effect, and we found increased expression per cell (blue color, Figure 5A) and/or more cells expressing each of these genes (filled circles, Figure 5A). We further investigated whether expression of selected genes was restored to sham levels, by looking at the normalized average gene expression in the capillary-1 cluster, which had the most dramatic changes related to expression of genes involved in vessel maturation and the maintenance of the blood-brain barrier. We selected 15 genes that were changed within the biological pathways of blood vessel maturation and maintenance of the blood-brain barrier. These genes were selected based on their relevance to the processes of cell-cell junction formation, pericyte recruitment and cytoskeleton assembly (Figure 5A). Of these 15 selected genes, expression levels of 10 of 15 were restored to Sham-α-IC levels after chronic treatment with α-VLA4, with 8 of 15 restored after chronic treatment with α-VCAM1. The remaining 5-7 genes were restored to ∼60% of their expression relative to levels in the Sham-α-IC cells after chronic antibody treatment. Notably, gene expression of *Ctnnb1* (Catenin B), a protein within adherens junctions, followed this pattern (Figure 5B). Cell-cell junctions are critical for communication between cells of the neurovascular unit and establishment of an intact blood-brain barrier, so we next investigated individual genes involved in pericyte recruitment, including growth factors and others that modulate the basement membrane and extracellular matrix. Expression of the growth factor *Edn3* (Endothelin 3; Figure 5C) and the collagenase *Plod2* (Procollagen Lysyl Hydroxylase 2; Figure 5D) were both reduced by stroke, and restored to sham levels after α-VCAM1 or α-VLA4 chronic treatment.

**Figure 5.**
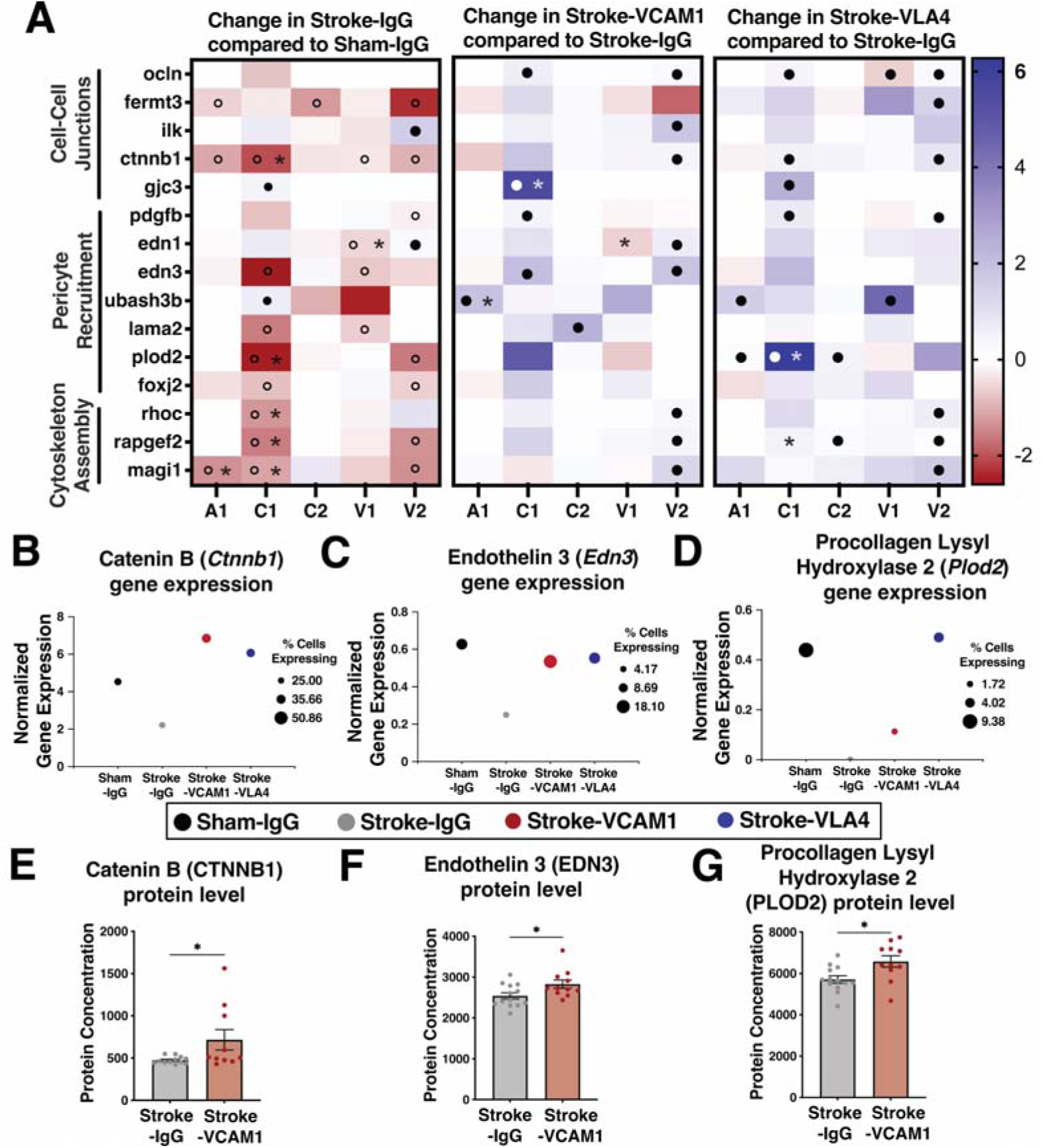
Genes involved in cell-cell junction formation, pericyte recruitment and cytoskeleton assembly are stimulated after anti-VCAM1 and anti-VLA4 treatment. A) Heat map comparing the Log_2_Fold Change of selected genes involved in cell-cell junction formation, pericyte recruitment or cytoskeleton from each endothelial cell cluster. Three different bioinformatic comparisons are depicted from left to right: Stroke-α-IC v. Sham-α-IC; Stroke-α-VCAM1 v. Stroke-α-IC; Stroke-α-VLA4 v Stroke-α-IC. A: Artery cell cluster; C1: Capillary-1 cluster; C2: Capillary-2 cluster; V1: Vein-1 cluster; V2: Vein-2 cluster. A star represents a significant Log_2_Fold Change (univariate *p* < 0.05), and a circle represents a change >10% of cells expressing the selected gene. Open circles represent a decrease in the percent of cells expressing that gene, while filled circles represent an increase. B-D) Normalized gene expression (scaled transcript counts and percent of population expressing) of selected genes involved in cell-cell junction formation (B; *Ctnnb1*), pericyte recruitment (C; *Edn3*) and blood-brain barrier maintenance (D; *Plod2*) in the capillary-1 cell cluster, and how this changes with treatment. Dot size represents the percent of cells within each cluster expressing the individual gene. E-G) Protein concentration from SomaLogic proteomics of Catenin B (E), Endothelin-3 (F), and Procollagen Lysyl Hydroxylase 2 (G). \**p* < 0.05, Mann-Whitney U test.

Next, we performed proteomics on plasma isolated from the Stroke-α-IC and Stroke-α-VCAM1 (Chronic) groups, utilizing the full SomaLogic Proteomics Panel (5,284 total aptamers that measure plasma proteins in mice). Chronic α-VCAM1 treatment also exhibited a vasculoprotective signature. Consistent with the transcriptomic data, we observed elevated plasma protein expression of Catenin B (CTNNB1; Figure 5E), Endothelin-3 (EDN3; Figure 5F), and Procollagen Lysyl Hydroxylase 2 (PLOD2; Figure 5G). We next utilized a *p* < 0.05 cutoff to identify all proteins significantly changed after chronic α-VCAM1 treatment. We observed moderate changes in additional proteins involved in vascular growth and maintenance. These include increases with chronic α-VCAM1 treatment in Thrombin, Prolyl-4 Hydroxylase Alpha-2, Tenascin-X, Leucine-rich Repeat-containing G Protein-coupled Receptor 5, Transmembrane Serine Protease 6, and GSK3B-interacting Protein, which are all involved in vascular growth and maintenance. Conversely, the concentration of proteins associated with blood-brain barrier dysfunction such as Collagen 8A1, Sarp-1, Endothelin-converting Enzyme-1, Collagen 11A2, and Matrix Metalloproteinase-16 decreased after chronic α-VCAM1 treatment (Supplementary Table 2). Additionally, proteins that are downstream of the VLA4/VCAM1 signaling axis that function to open the blood-brain barrier, such as Spastin and Rho Guanine Nucleotide Exchange Factor 2, were also decreased by chronic α-VCAM1 treatment.

Given the strong biological signature of vessel growth and maturation observed with both single cell RNA sequencing and plasma proteomics, we next used immunostaining to formally quantify these processes (Figure 6). We first analyzed the number and length of CD31+ blood vessels in the peri-infarct cortex to assess vessel growth. There was no difference in total vessel number (Supplementary Figure 8), but α-VCAM1 treated mice demonstrated a trend towards longer vessels (Figure 6A) and chronic α-VLA4 treated mice had longer vessels than isotype control antibody treated mice after stroke (Figure 6D). We next analyzed vascular pericyte coverage as a measure of blood vessel maturation and blood-brain barrier structural integrity. Both chronic α-VCAM1 and α-VLA4 treatment increase CD13+ pericyte coverage of CD31+ blood vessels by 73% and 68%, respectively, in the peri-infarct cortex (Figure 6B,C,E,F). In the contralateral cortex, there was a trend towards increased coverage in α-VCAM1 treated mice (Figure 6C), while chronic α-VLA4 treatment increased pericyte coverage by 68% (Figure 7F). Finally, due to the critical role of pericytes in both forming and maintaining an intact blood-brain barrier, we analyzed extravascular fibrinogen leakage into the brain parenchyma. Chronic α-VCAM1 treatment completely eliminated extravascular fibrinogen leakage in the peri-infarct cortex, and there was also a strong trend towards reduced leakage in the stroke scar (Figure 7A,B). Similarly, chronic α-VLA4 significantly reduced extravascular fibrinogen leakage in both the stroke scar and peri-infarct cortex compared to Stroke-α-IC mice (Figure 7C,D). Thus, chronic treatment with α-VCAM1 and α-VLA4 preserved cerebrovascular structural and functional integrity in chronic stroke.

**Figure 6.**
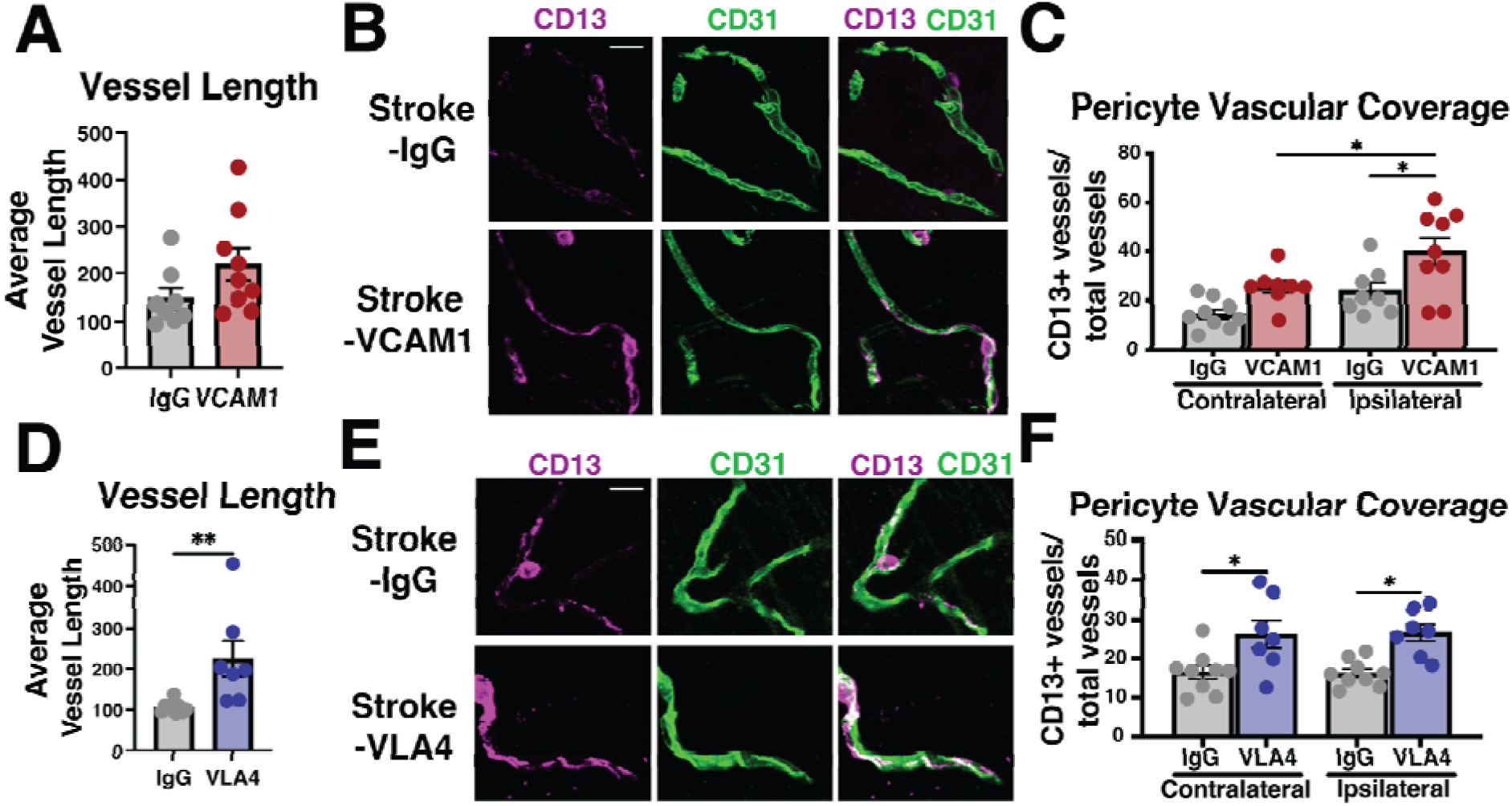
Effects of anti-VCAM1 and anti-VLA4 chronic treatment on vascular length and coverage by pericytes. A) Total vessel length in the peri-infarct cortex after 3 weeks of chronic isotype control (Stroke-α-IC) or anti-VCAM1 (Stroke-α-VCAM1) antibody treatment, using endothelial staining for CD31 to define vasculature. B) Representative confocal images and (C) quantification of pericyte (CD13+) coverage of CD31+ vasculature in the peri-infarct cortex or contralateral cortical region of stroked mice treated with isotype control (Stroke-α-IC) or anti-VCAM1 (Stroke-α-VCAM1) antibody. D) Total vessel length in the peri-infarct cortex after 3 weeks of chronic isotype control (Stroke-α-IC) or anti-VLA4 (Stroke-α-VLA4) antibody treatment. E) Representative confocal images and (F) quantification of pericyte (CD13+) coverage of CD31+ vasculature in the peri-infarct cortex or contralateral cortical region of stroked mice treated with isotype control (Stroke-α-IC) or anti-VLA4 (Stroke-α-VLA4) antibody. Scale bar, 20uM; Statistics, Student’s t-test or 2-way ANOVA with Tukey’s post hoc test for group comparisons, Error bars, mean ± SEM; *p<0.05; **p<0.01.

**Figure 7.**
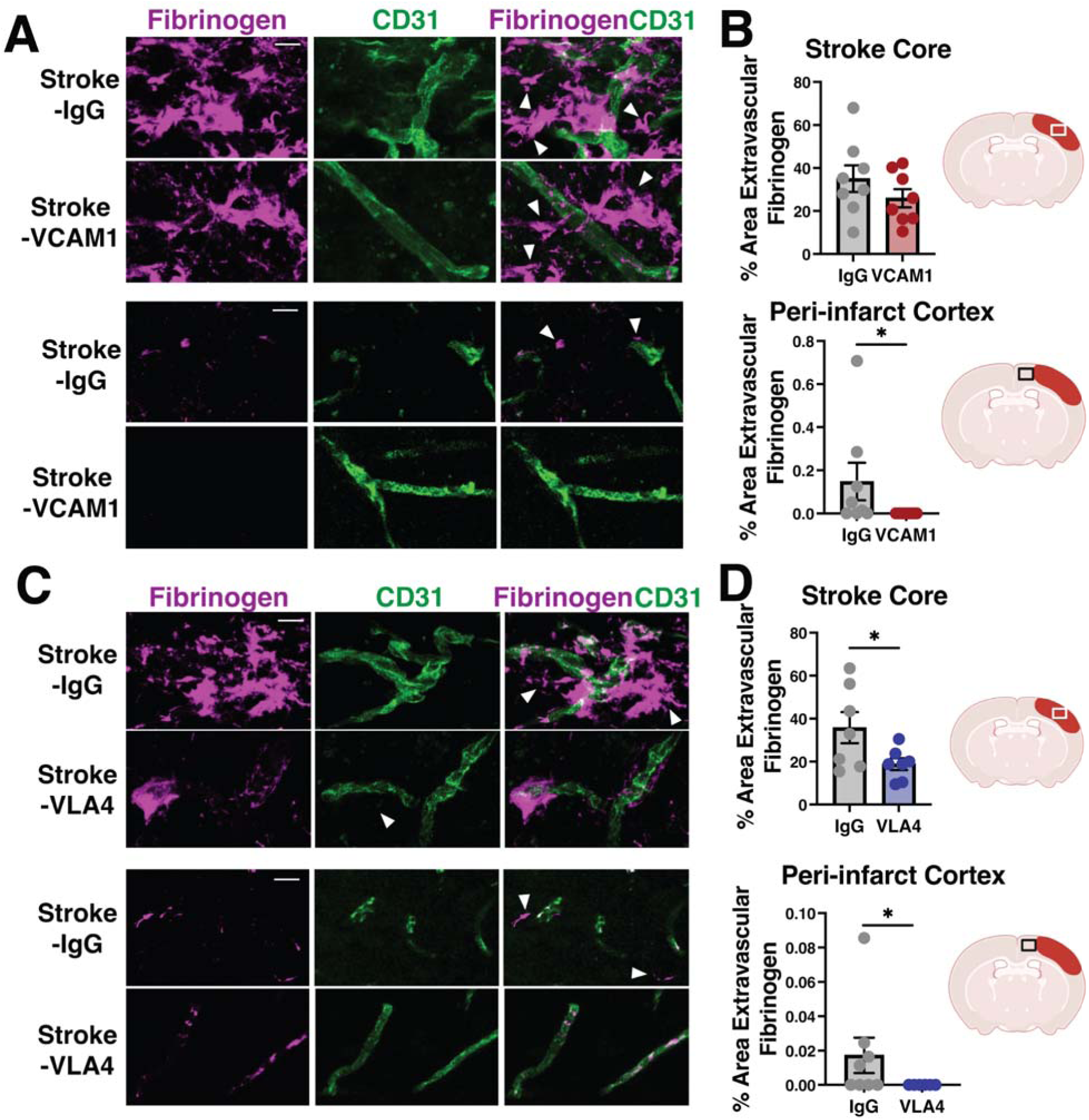
Anti-VCAM1 and anti-VLA4 chronic treatment reduces blood-brain barrier leakage. A) Representative confocal images and B) quantification of the percent area covered by extravascular fibrinogen in the stroke core (top) and peri-infarct cortex (bottom) of mice treated with chronic isotype control (Stroke-α-IC) or anti-VCAM1 (Stroke-α-VCAM1) antibody for 3 weeks. C) Representative confocal images and D) quantification of the percent area covered by extravascular fibrinogen in the stroke core (top) and peri-infarct cortex (bottom) of mice treated with chronic isotype control (Stroke-α-IC) or anti-VLA4 (Stroke-α-VLA4) antibody for 3 weeks. White arrows point to extravascular fibrinogen. Scale bar, 20 µM; Statistics, Student’s *t* test; Error bars, mean ± SEM; *p < 0.05.

## Discussion

We report here that a mouse model of infarct-induced neurodegeneration is characterized by chronic structural and functional blood-brain barrier deficits in the stroke scar and peri-stroke cortex. We observe loss of pericyte coverage and tight junction proteins on vasculature, as well as leakage of fibrinogen into the brain parenchyma. While immune cell transcriptional changes had largely resolved by ten weeks after stroke, endothelial cells retained a signal of chronic dysfunction. The reduction in vascular pericyte coverage and ZO-1 tight junctions at the protein level was supported by a reduction in expression of genes involved in tight junction formation and pericyte recruitment and retention. Additionally, we observed increased expression of genes related to blood-brain barrier permeability such as collagenases and matrix metalloproteinases. Venous endothelial cells also exhibited evidence of increased permeability, but rather than overt changes in gene expression among the population, the population of venous endothelial cells expressing endothelial permeability genes such as *Ctnnb1* and *Magi1* increased. The percent of venous endothelial cells expressing proliferation and stress response genes also increased, perhaps reflecting generally more porous and activated venous endothelium. Thus, there was an overall loss of vascular integrity and increased proliferation and stress response throughout the vascular bed observed via both immunostaining and single cell RNA sequencing. We hypothesized that cognitive decline late after stroke was dependent on this blood-brain barrier dysfunction, and then showed that we can prevent blood-brain barrier dysfunction and improve cognition by blocking the interaction between VLA4 and VCAM1 in mice of both sexes.

Our data is robust in terms of both demonstrating the clinical benefits and the biological mechanism by which the VCAM1 and VLA4 interaction acts on endothelial cells. To demonstrate cognitive benefit, we utilized two different antibodies to block the VLA4/VCAM1 axis, one against VCAM1 and one against VLA4. Chronic blockade with either blocking antibody robustly prevented infarct-induced neurodegeneration in both male and female mice. We observed this result in 3 experiments with anti-VCAM1 antibodies, twice in males and once in females. We then confirmed the protective effects of blocking this axis with an anti-VLA4 antibody in two additional experiments, one in males and one in females. In addition, we utilized two distinct cognitive tasks to interrogate memory (novel object recognition) and learning/executive function (Barnes maze) and observed improved cognitive function in each experiment. Using high-depth single cell RNA sequencing of endothelial cells from mice with chronic stroke, we saw reduced expression of genes associated with angiogenesis and maturation of the blood-brain barrier in mice with chronic stroke that were treated with isotype control antibody and developed cognitive deficits. Both chronic antibody treatments ameliorated these chronic stroke-induced gene expression changes in endothelial cells, pointing to preservation of normal cerebrovasculature as a primary mechanism of action. Plasma proteomics from mice treated with chronic anti-VCAM1 compared to isotype control antibody also demonstrated comparable plasma protein changes. And finally, chronic treatment with either antibody improved vascular pericyte coverage and ameliorated blood-brain barrier leakiness.

This new work refines our model of the mechanism of infarct-induced neurodegeneration (Figure 8). In infarct-induced neurodegeneration we observe a chronic inflammatory process where B and T lymphocytes are activated and traffic to the brain in the weeks after stroke via the VLA4/VCAM1 interaction.^13^ This requires and also further stimulates persistent vascular dysfunction which is characterized by a disrupted blood-brain barrier that is leaky to pathological molecules such as fibrinogen. In turn, the leaky blood-brain barrier also stimulates neuroinflammation via activated endothelium that upregulates VCAM1 expression, triggering a positive feedback loop that continues to propagate chronic inflammation and blood-brain barrier dysfunction. Over time, both blood-brain barrier leakage and immune cell infiltration could lead to cognitive decline, and as discussed below, our data suggests that blood-brain barrier leakage is likely more important (Figure 8, left). When we block the VLA4/VCAM1 axis via chronic treatment with anti-VCAM1 or anti-VLA4, we disrupt the positive feedback loop by promoting generation of mature blood vessels with intact tight junctions, restoring pericytes around the vessels and producing a mature and intact blood-brain barrier (Figure 8, right). This strongly supports the hypothesis that vascular integrity is the primary mechanism by which chronic anti-VCAM1 and anti-VLA4 treatment preserve cognitive function.

**Figure 8.**
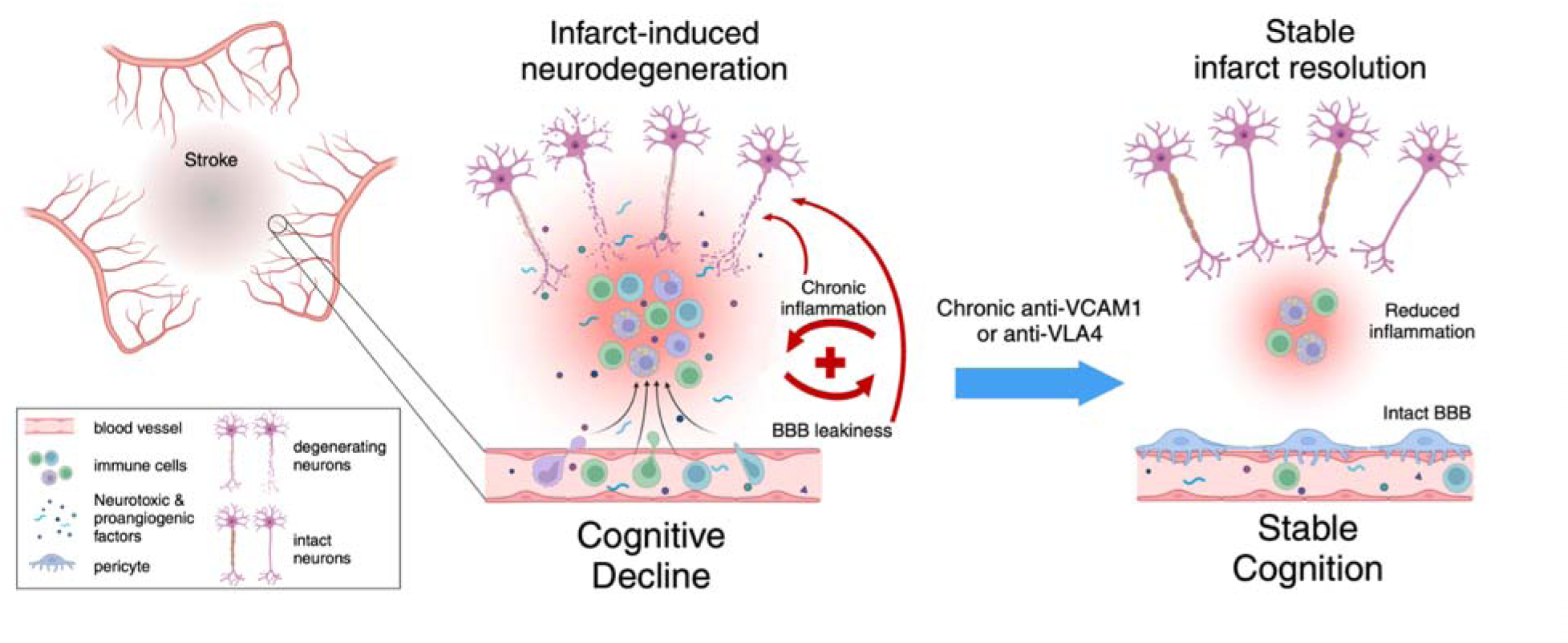
Proposed mechanism of the VLA4/VCAM1 axis in preventing infarct-induced neurodegeneration. The diagram depicts our proposed hypothesis of how infarct-induced neurodegeneration, a chronic process, can be prevented with chronic blockade of the VLA4/VCAM1 axis. We propose that infarct-induced neurodegeneration occurs when the initial inflammation due to stroke fails to resolve, resulting in persistent vascular dysfunction, chronic immune cell trafficking to the brain, chronic blood brain barrier leakiness and subsequent cognitive decline. Chronic blockade of the VLA4/VCAM1 signaling axis restores blood brain barrier integrity, reduces immune cell trafficking to the brain, and prevents chronic cognitive decline after stroke. Diagram created using BioRender.

This work represents the first time that VCAM1 and VLA4 have been blocked chronically after stroke, however there is precedent that blockade of this signaling pathway might affect both immune cell infiltration and blood-brain barrier integrity. In rodent models of stroke, acute dosing of anti-VLA4 robustly reduces leukocyte infiltration into the brain, and also reduces acute infarct size in some but not all studies.^39–43^ In rodent models of the autoimmune disease multiple sclerosis, experimental autoimmune encephalitis, anti-VLA4 treatment reduces B and T lymphocyte trafficking into the brain, demyelinating lesion size and axonal loss.^44–47^ Additionally, while there is minimal leukocyte trafficking to the brain during normal aging, chronic anti-VCAM1 treatment of aged mice reduces neuroinflammation and endothelial cell activation, and restores cognitive function.^23^

Our study also highlights the effects of the VLA4/VCAM1 interaction on vascular permeability. These effects are consistent with prior knowledge of the VLA4/VCAM1 axis. Opening the blood-brain barrier is necessary for leukocyte diapedesis into the brain. Indeed, the VLA4/VCAM1 axis has a direct effect on vascular permeability in cancer models, where activating this axis induces hyperpermeability of an endothelial monolayer *in vitro*.^48,49^ This increased permeability is induced by structural rearrangement of tight and adherens junctions, as well as cytoskeletal reassembly, mediated via multiple genes including *Ctnnb1*, *Rhoc* and *Magi1* which are all downstream of VCAM1 signaling in endothelial cells.^50^

Despite significant pro-inflammatory activation of immune cells acutely,^51,52^ we observed very few gene changes 10 weeks after stroke. While it is possible that subtle gene expression changes may be masked due to processing artifacts during tissue digestion, we did observe a robust increase in the proportion of peripheral immune cells in the brain after stroke.^53^ And we do observe high numbers of B and T lymphocytes in the brain after stroke, which were significantly reduced after anti-VCAM1 treatment. This reduction in B and T lymphocytes was also associated with improved cognition in our study. This is consistent with our previous study in adult mice where we showed that ablating B cells beginning 5 days after stroke prevents delayed cognitive decline.^13^ We also demonstrated that humans who die with infarcts and dementia have increased numbers of B cells in and around the stroke, and higher MBP antibody titers are associated with worse cognitive trajectories in stroke survivors,^13,54,55^ suggesting this mechanism is likely important in humans with chronic stroke. However, in those prior studies, neither high B lymphocyte numbers in infarcts nor high anti-MBP antibody titers were present in all people with infarcts and dementia or cognitive decline after stroke. This implies that some other factor may be more important in humans with infarct-induced neurodegeneration.

In line with this concept, it was unexpected and intriguing that anti-VLA4 treatment had a strong positive effect on cognition without meaningfully reducing B lymphocyte numbers in the brain in our histology analysis. Similarly, the effects of both antibodies on immune cell gene expression were underwhelming. Thus, while B cells and/or autoimmune responses may be relevant for some and/or required early after stroke to instigate infarct-induced neurodegeneration, they may not be required in the brain for perpetuation of infarct-induced neurodegeneration in people, and it may rather be changes in blood-brain barrier function that drive longer term cognitive changes.

In further support of a role for cerebrovascular defects driving infarct-induced neurodegeneration, we saw dramatic stroke-induced changes in gene expression in endothelial cells 10 weeks after stroke. High depth sequencing allowed a high-resolution look at how different vessel segments respond to stroke. We found that endothelial cells from the arterial segment exhibited the most changes in total gene expression between mice from the sham and stroke groups, while venous endothelial cells changed the least. This is consistent with another single cell RNA sequencing study investigating how endothelial cells respond to age or acute LPS injection,^56^ where venous cells respond less than other vessel segments. However, in that study, capillary rather than arterial endothelial cells exhibited the most transcriptional change. This difference may reflect arterialization of new vessels that grow in the recovery phase after stroke. This has been observed after ligation of the middle cerebral artery in rats, where both the diameter and lengths of collaterals on the brain surface significantly increase one month later.^57^

Our findings that elevated VCAM1 activity is associated with decreased pericyte coverage and fibrinogen leakage across the blood-brain barrier in chronic stroke are detrimental in infarct-induced neurodegeneration are consistent with findings in other neurodegenerative diseases. Chronic blood-brain barrier leakage is associated with cognitive decline in both aging and Alzheimer’s disease.^18,58,59^ Moreover, chronic blood-brain barrier leakage itself can directly contribute to cognitive decline via fibrinogen, which contributes to neuroinflammation and cognitive decline in rodent models of brain injury, Alzheimer’s disease, and multiple sclerosis.^16,60–62^ This may be mediated via increased synapse elimination or neuronal senescence resulting in cognitive dysfunction, which has been causally linked to neuroinflammation and blood-brain barrier breakdown after stroke.^63,64^ Although it is well known that there is acute pericyte loss in the infarct core after stroke,^65–67^ as well as acute blood-brain barrier leakiness,^15,68,69^ little is known about vascular integrity in the chronic phase after stroke. Here, we have demonstrated for the first time, to our knowledge, that vascular dysfunction persists chronically for months after stroke, and is characterized by loss of pericyte coverage, reduced ZO-1 tight junction levels and increased extravascular fibrinogen leakage. Notably, pericyte vascular coverage is reduced in rodent models of aging, and there is deficient pericyte remodeling after injury.^70,71^ Aging is also a major risk factor for both stroke and dementia, suggesting that impaired vascular integrity with age may be an underlying causal mechanism. New work in our laboratory also demonstrates a prominent loss of pericytes in infarct-induced neurodegeneration in people, with a median value close to zero pericyte coverage of vessels in people who died with infarcts and dementia, but not those who died with infarcts and no dementia (unpublished). We show here that this also happens in the mouse model, and excitingly that chronic anti-VCAM1 and anti-VLA4 treatment reduce this leakage and prevent cognitive decline in chronic stroke. Plasma sVCAM1 levels are also high in people with other types of dementia including vascular dementia, Parkinson’s disease and Alzheimer’s disease, where blood-brain barrier leakiness may precede cognitive decline.^21,35,36,38,72^ Together, this evidence supports the VLA4/VCAM1 axis as a strong translational therapeutic target to improve vascular integrity and preserve cognition in neurodegenerative disease.

Anti-VLA4 antibody therapy with Natalizumab is FDA approved and likely will help prevent infarct-induced neurodegeneration in people. It is used routinely in multiple sclerosis where it prevents blood-brain barrier leakiness and promotes better outcomes.^73–75^ Indeed, there is some evidence that Natalizumab improves cognitive outcomes in multiple sclerosis.^76,77^ In acute stroke, it is proven safe but had no impact on infarct size or 90 day cognitive outcomes.^78,79^ However, in these clinical trials (ACTION I and ACTION II), a single acute dose of Natalizumab was given between 9 and 24 hours after stroke. Here, when we used a single dose in mice at 4 hours after stroke, this also did not improve cognition. However, when we treat mice chronically, beginning on day 4, we see robust prevention of infarct-induced neurodegeneration. Thus, chronic therapy regimens, similar to those used in treatment of multiple sclerosis, may be necessary to prevent cognitive decline after stroke.

In conclusion, we report that vascular dysfunction persists chronically after stroke, and that blocking the VLA4/VCAM1 axis with chronic antibody treatment preserves vascular integrity and prevents cognitive decline in a middle-aged mouse model of infarct-induced neurodegeneration. We used single cell RNA sequencing, plasma proteomics, and immunostaining to demonstrate that both anti-VCAM1 and anti-VLA4 promote cerebrovascular health in chronic stroke. They both increase structural integrity as represented by vascular pericyte coverage and restore the blood-brain barrier chronically after stroke. Future work is needed to further test these strategies to restore pericyte coverage and restore the blood-brain barrier in chronic stroke and determine whether these mechanisms will benefit people with infarct-induced neurodegeneration.

## Data availability

All data is available upon request from the corresponding author. Single cell RNA sequencing data can be accessed through GEO (Accession # GSE300564).

## Supplementary Material

Supplementary material is available at *Brain* online.

## Supplementary Data

**Supplementary Figure 1.**
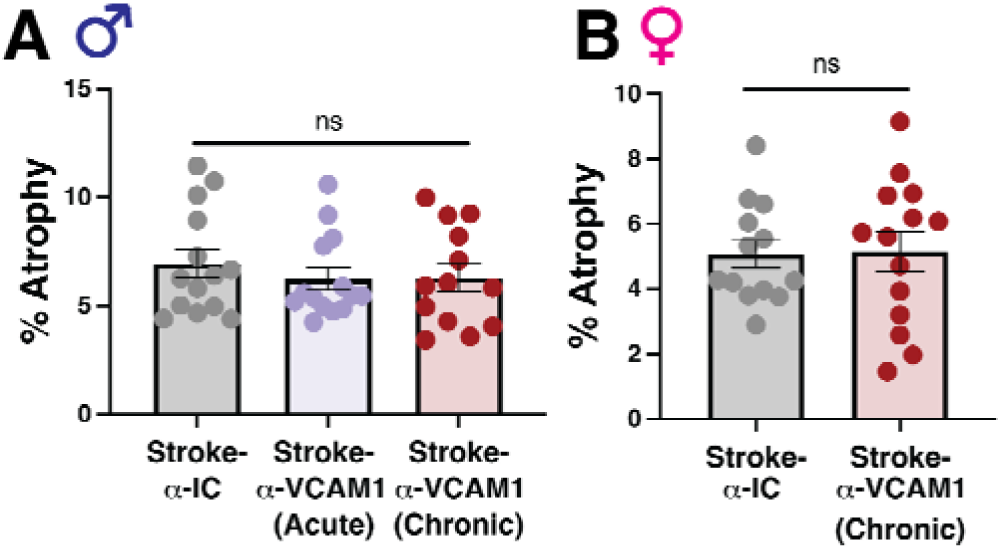
Chronic brain atrophy is unchanged by anti-VCAM1 treatment in middle-aged mice. Quantification of total brain atrophy in 10 month old male (A) or female (B) mice. Mice were treated with isotype control (Stroke-α-IC; n=14 male, n=13 female) or anti-VCAM1 (Stroke-α-VCAM1) antibody acutely (n=14 male) or chronically (n=13 male, n=14 female). Statistics, Student’s *t* test or one-way ANOVA with Tukey’s post-hoc test; Error bars, mean ± SEM.

**Supplementary Figure 2.**
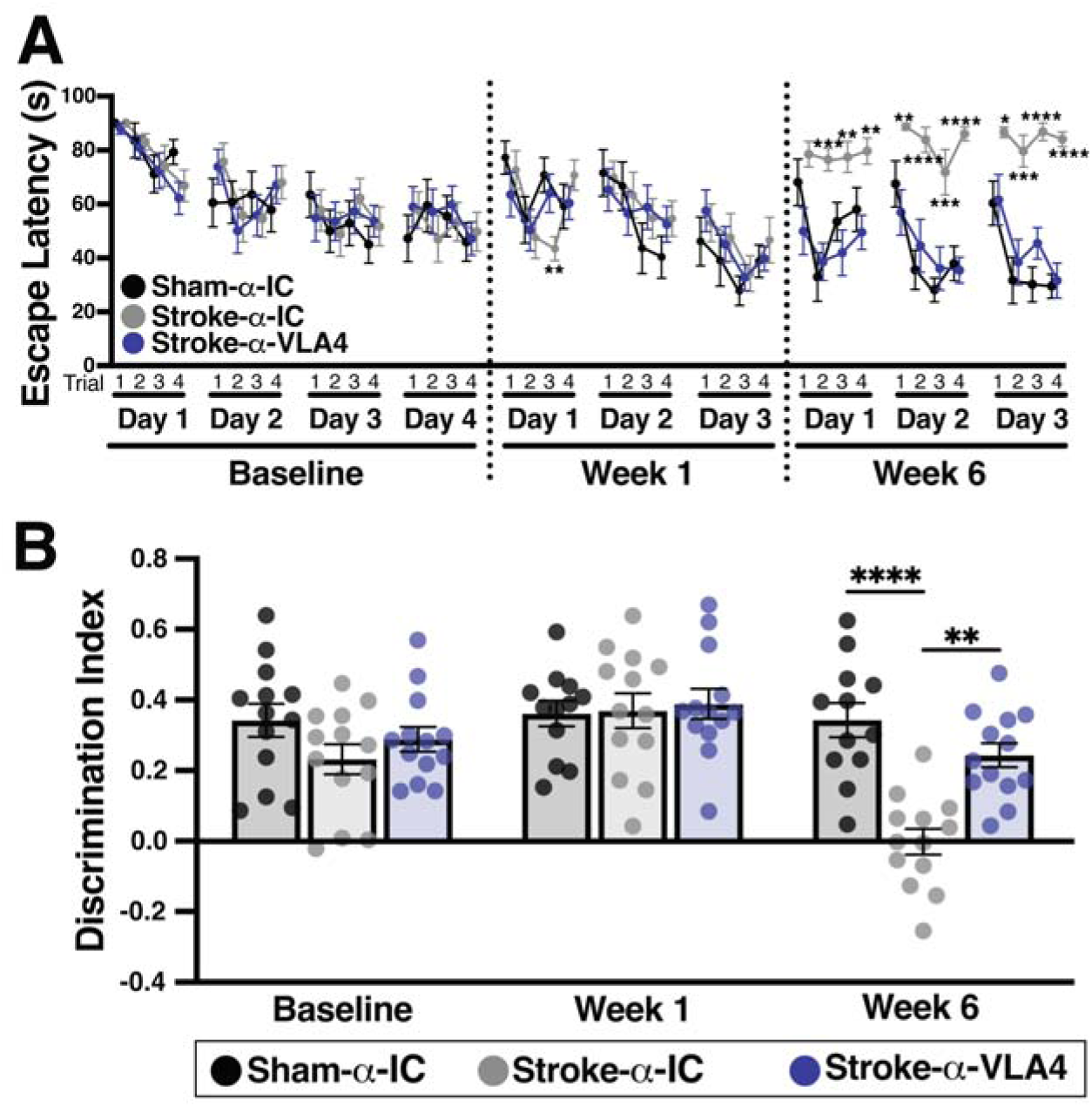
Anti-VLA4 chronic antibody treatment prevents cognitive decline in middle-aged female mice. A) Escape latency on the Barnes maze and B) discrimination index on the novel object recognition task for 10 month old female mice treated with isotype control antibody (α-IC; n=13 sham and 13 stroke or anti-VLA4 (α-VLA4; n=13) chronically. Statistics, 2-way repeated measures ANOVA with Bonferroni’s post-hoc test; Error bars, mean ± SEM; *p < 0.05; **p < 0.01; ***p < 0.001; ****p < 0.0001.

**Supplementary Figure 3.**
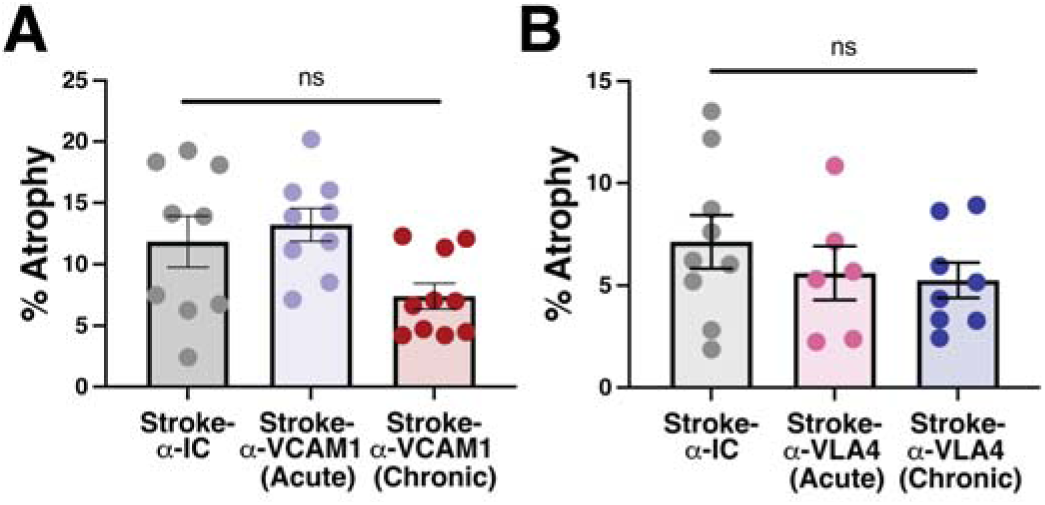
Chronic brain atrophy is unchanged by anti-VCAM1 or anti-VLA4 treatment in adult mice. Quantification of total brain atrophy in 3 month old male mice. In (A), mice were treated with isotype control (Stroke-α-IC; n=9) or anti-VCAM1 (Stroke-α-VCAM1) antibody acutely (n=9) or chronically (n=10) for 3 weeks. In (B) mice were treated with isotype control (Stroke-α-IC; n=9) or anti-VLA4 (Stroke-α-VLA4) antibody acutely (n=6) or chronically (n=8). Statistics, one-way ANOVA with Tukey’s post-hoc test; Error bars, mean ± SEM.

**Supplementary Figure 4.**
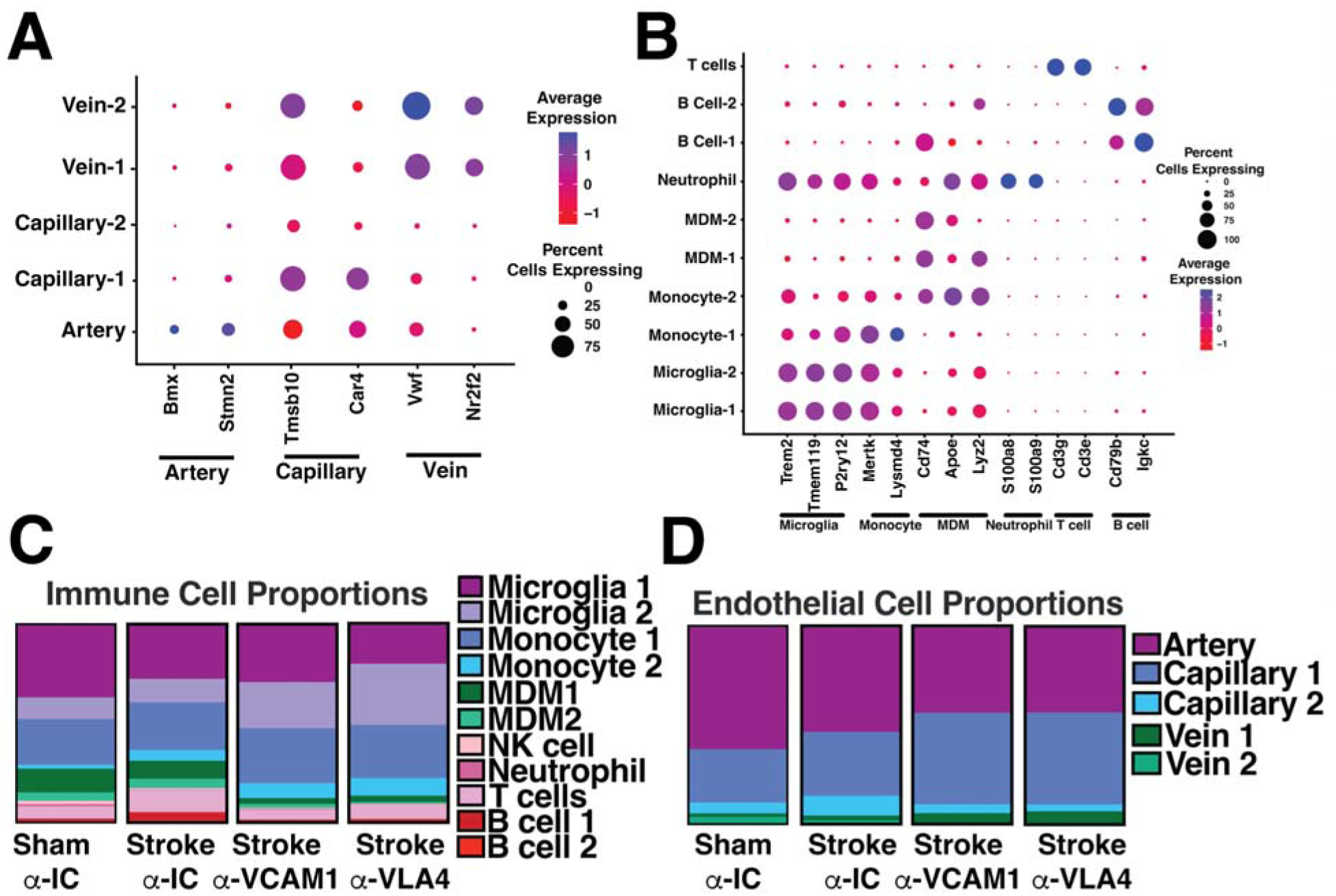
Chronic anti-VCAM1 and anti-VLA4 treatment does not alter immune cell phenotype chronically after stroke. A dotplot of cell marker genes was used to annotate cell clusters in (A) endothelial and (B) immune cells. A graphical representation of the proportion of cells collected from each cluster, separated by sample for immune (C) and endothelial (D) cells.

**Supplementary Figure 5.**
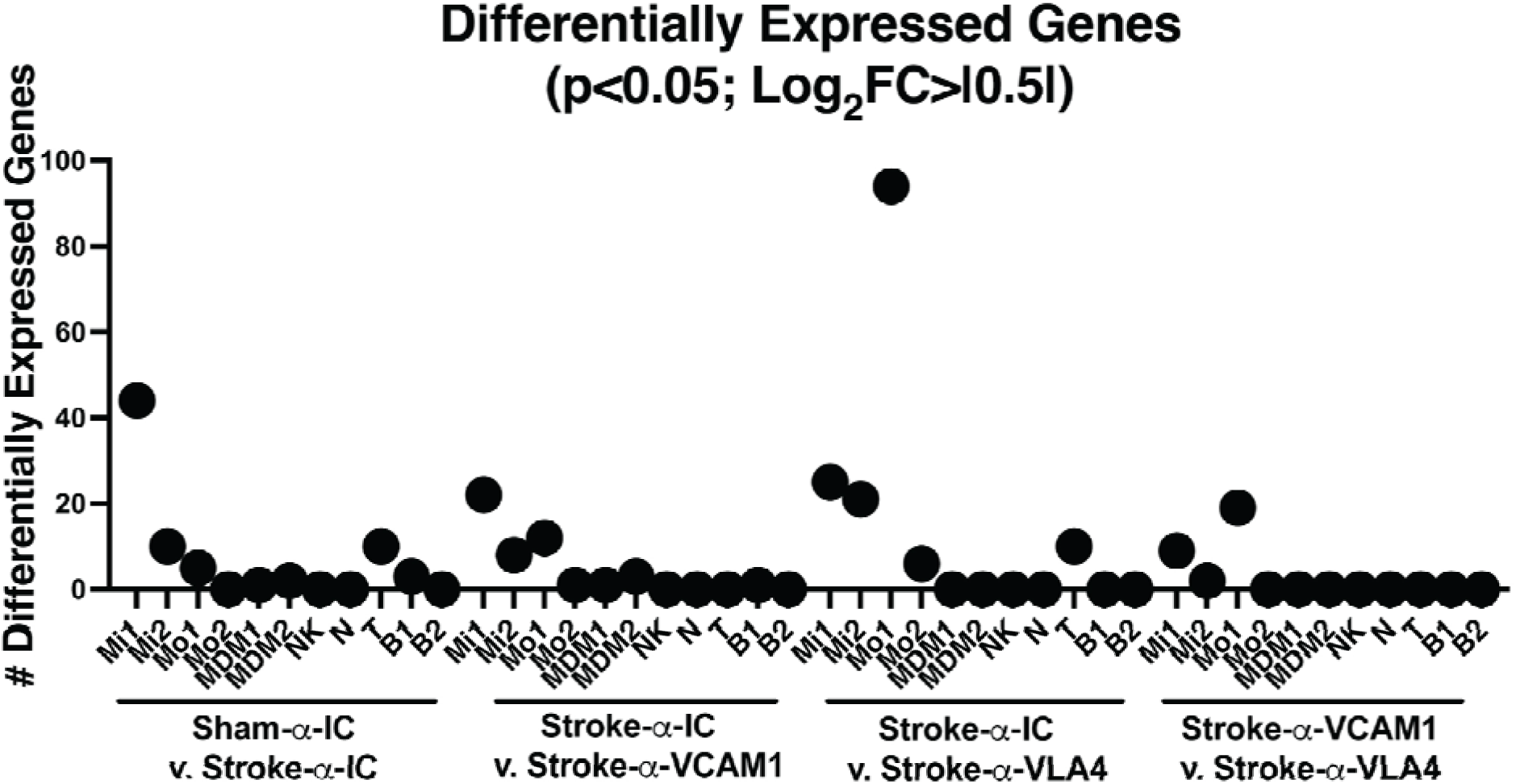
A graphical representation of the number of differentially expressed genes in immune cell subpopulations defined by p<0.05 and Log_2_FoldChange>|0.5|. Mi, Microglia; Mo, Monocyte; MDM, Monocyte-derived macrophage; NK, Natural Killer cell; N, Neutrophil; T, T cell; B, B cell.

**Supplementary Figure 6.**
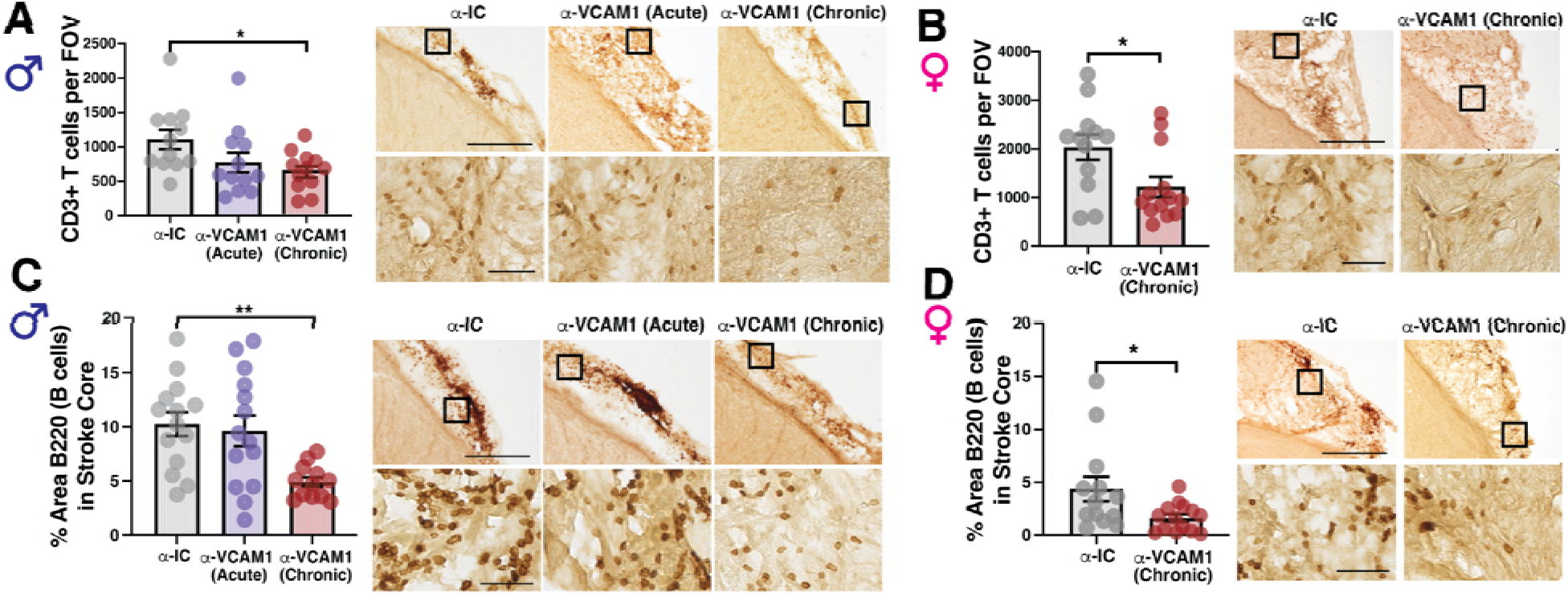
A) Quantification of CD3+ T cell infiltration per field of view (FOV) into the stroke core in male and B) female mice, 8 weeks after stroke. C) Percent area covered by B220+ (B cell) immunostaining in the stroke core of male and D) female mice, 8 weeks after stroke. Statistics, one-way ANOVA with Tukey’s post hoc test (A,C), or two-tailed Student’s *t* test (B,D). Scale bar, 300μM (top row) or 50μM (bottom row); Error bars, mean ± SEM; *p<0.05; **p<0.01.

**Supplementary Figure 7.**
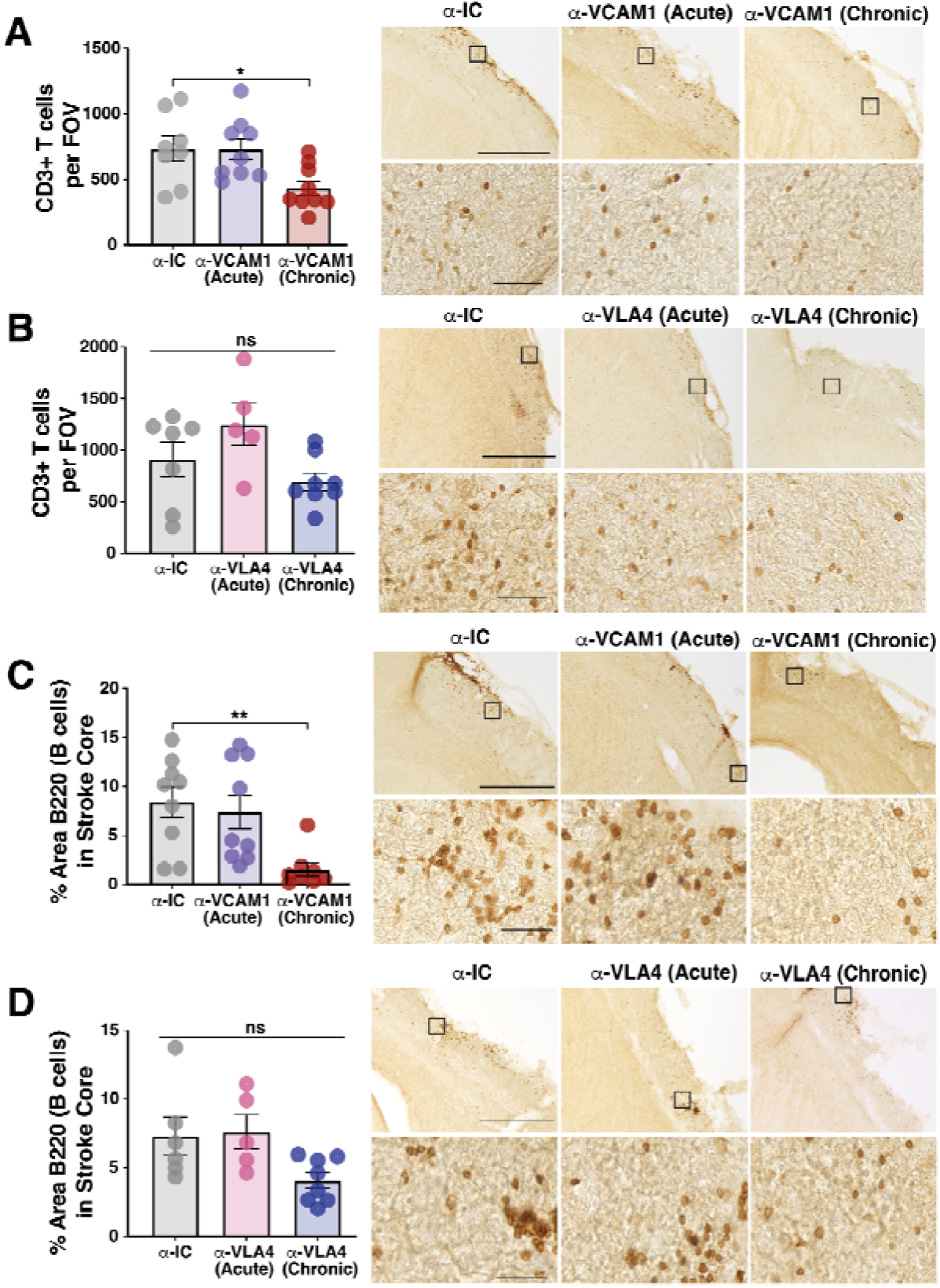
Chronic anti-VCAM1 is more effective at reducing lymphocyte trafficking than chronic anti-VLA4 treatment. A) Quantification and representative bright field images of CD3+ T cells in the stroke core after isotype control (α-IC; n=8), anti-VCAM1 acute (Stroke-α-VCAM1-Acute; n=9) or chronic (Stroke-α-VCAM1-Chronic; n=9) treatment, or B) isotype control (α-IC; n=7), anti-VLA4 acute (Stroke-α-VLA4-Acute; n=5) or chronic (Stroke-α-VLA4-Chronic; n=8) antibody treatment 3 weeks after stroke. C-D) Quantification and representative bright field images of B220+ B cells in the stroke core in the same mice. Scale bar, 300μM (top row) or 50μM (bottom row); statistics, 2-way ANOVA with Tukey’s post hoc test for group comparisons; Error bars, mean ± SEM; *p<0.05; **p<0.01; ***p<0.001; ****p<0.0001.

**Supplementary Figure 8.**
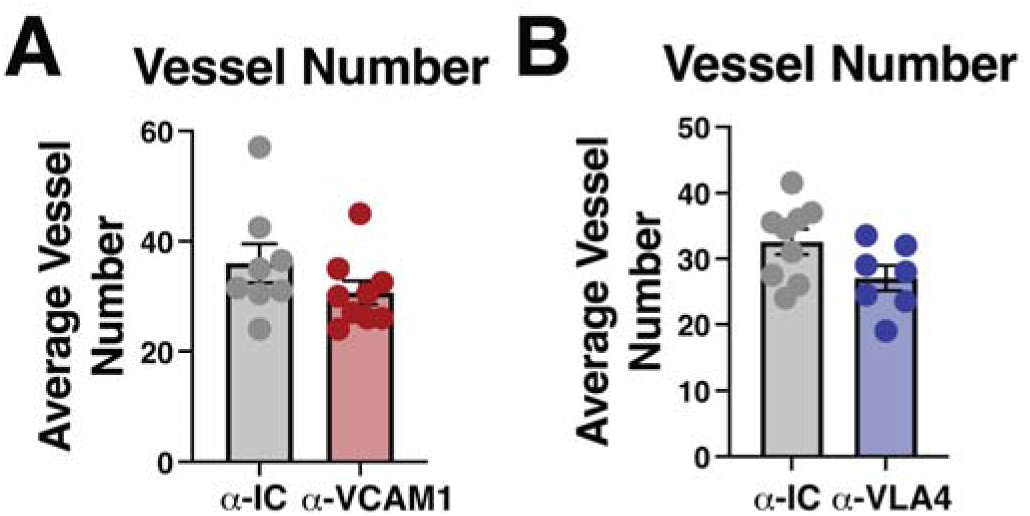
Total vessel number is unchanged after chronic anti-VCAM1 or anti-VLA4 treatment. Quantification of total blood vessel number in 3 month old male mice. In (A), mice were treated with isotype control (α-IC, n=8) or anti-VCAM1 (α-VCAM1) antibody (n=9) every 3 days for 3 weeks. In (B) mice were treated with isotype control (α-IC, n=9) or anti-VLA4 (α-VLA4, n=7) antibody every 3 days for 3 weeks. Statistics, Student’s *t* test; Error bars, mean ± SEM.

**Supplementary Table 1.** Cell markers used to annotate clusters isolated during single cell RNA sequencing of CD45+ and CD31+ cells. We isolated CD31+ endothelial and CD45+ immune cells from the stroke scar of 10 month old male C57BL/6J mice, 10 weeks after stroke or sham surgery. We used the UMAP algorithm to independently cluster each cell type by similarity, and utilized cell-specific markers to annotate the resulting clusters.

| Dataset | Cell Sub-type | Marker | Reference(s)Figure 1B |
| --- | --- | --- | --- |
| CD31 | Artery | *Stmn2* | 56,80 |
|  |  | *Bmx* | 56,80 |
| CD31 | Capillary | *Tmsb10* | 56,80 |
|  |  | *Car4* | 56,80 |
| CD31 | Vein | *Vwf* | 56,80 |
|  |  | *Nr2f2* | 56,80 |
| CD45 | Microglia | *Tmem119* | 81 |
|  |  | *Trem2* | 81 |
|  |  | *P2ry12* | 81 |
| CD45 | Monocyte | *Mertk* | 82 |
|  |  | *Lysmd4* | 82 |
| CD45 | Monocyte-derived Macrophage | *Apoe* | 82 |
|  |  | *Lyz2* | 82 |
|  |  | *Cd74* | 82 |
| CD45 | Neutrophil | *S100a8* | 83 |
|  |  | *S100a9* | 83 |
| CD45 | T cell | *Cd3g* | 82 |
|  |  | *Cd3e* | 82 |
| CD45 | B cell | *Igkc* | 83 |
|  |  | *Cd79b* | 83 |

**Supplementary Table 2.** Plasma protein concentrations from SomaLogic Proteomics. We measured the concentration of 5,284 proteins from the plasma with SomaLogic Proteomics. Mice were 10 month old males and treated with isotope control or anti-VCAM1 antibody chronically until 10 weeks after stroke. We used a Mann Whitney U Test to determine whether protein concentrations were altered by antibody treatment. A univariate Spearman correlation was performed to analyze the relationship between protein concentration, regardless of treatment group, and cognitive function. To assess cognitive function, a cognitive score was calculated by averaging the Barnes Maze performance from trials 3&4 of the final day of testing for each mouse. Here, a lower cognitive score represents better cognitive function.

| Protein Symbol | Protein Name | Protein Concentrations |  | Correlation with Barnes Maze performance |  |
| --- | --- | --- | --- | --- | --- |
|  |  | IgG: VCAM1 Ratio | Mann Whitney P value | Spearman rho | Spearman P value |

